# Effective Dynamics of Nucleosome Configurations at the Yeast *PHO5* Promoter

**DOI:** 10.1101/2020.05.02.072470

**Authors:** Michael Roland Wolff, Andrea Schmid, Philipp Korber, Ulrich Gerland

## Abstract

Chromatin dynamics are mediated by remodeling enzymes and play crucial roles in gene regulation, as established in a paradigmatic model, the yeast *PHO5* promoter. However, effective nucleosome dynamics, i.e. trajectories of promoter nucleosome configurations, remain elusive. Here, we infer such dynamics from the integration of published single-molecule data that capture multi-nucleosome configurations for repressed to fully active *PHO5* promoter states with other existing histone turnover and new chromatin accessibility data. We devised and systematically investigated a new class of “regulated on-off-slide” models simulating global and local nucleosome (dis)assembly and sliding. Only seven of 68 145 models agreed well with all data. All seven models involve sliding and the known central role of the N-2 nucleosome, but regulate promoter state transitions by modulating just one assembly rather than disassembly process. This is consistent with but challenges common interpretations of previous observations at the *PHO5* promoter and suggests chromatin opening by binding competitions.

## Introduction

Eukaryotic DNA is packaged inside the nucleus with several layers of compaction. The most basic layer consists of nucleosomes, where 147 bp of DNA are wrapped around a histone protein octamer (Luger et al., 1997). Nucleosomes occupy most of the genome and hinder binding of other proteins to DNA, for example transcription factors (Venter et al., 1994; Bell et al., 2011; Lai and Pugh, 2017), replication machinery (Chang et al., 2016) and DNA repair enzymes (Hauer and Gasser, 2017). This hindrance can be overcome by chromatin remodeling enzymes. Such “remodelers” bind to nucleosomes and convert the energy from ATP hydrolysis into sliding, ejection, (re-)assembly or restructuring of nucleosomes (Bartholomew, 2014; Zhou et al., 2016; Clapier et al., 2017). Hence, nucleosomes are not static, but are constantly replaced to varying degrees throughout the genome by remodeling processes, e.g., in the context of transcription (Dion et al., 2007). To decipher the resulting nucleosome dynamics, detailed measurements of both, steady state and dynamic quantities are key.

Nucleosome occupancy, a steady-state quantity, is measured in a cell-averaged way with different techniques (reviewed in Lieleg et al., 2015). For example, non-nucleosomal DNA is digested with an endonuclease, like MNase, followed by high-throughput sequencing of the nucleosome-protected DNA. Or, DNA is cleaved close to the nucleosome dyad (= midpoint) position by hydroxyl radicals generated *in situ* at cysteine residues artificially introduced into histones, and the resulting DNA fragments are sequenced (Brogaard et al., 2012). After paired-end sequencing, the latter method also gives the distances between neighboring nucleosomes. While such data sets provide quite reliable information on single individual nucleosome positions and on nucleosome occupancy on population average, they have two disadvantages: First, the nucleosome occupancy in absolute terms, i.e., the fraction of molecules where a certain position falls within a nucleosome, cannot be determined as only nucleosomal DNA was scored and non-nucleosomal DNA lost from the analysis (for detailed discussion and solution to this question see Oberbeckmann et al., 2019). Second, the information on the interactions or coupling between several nucleosomes along the same DNA molecule in genomic regions of interest is lost, i.e., the obtained occupancy map corresponds only to the one-particle density.

These two disadvantages prevent a more detailed modeling of nucleosome dynamics. In this study, we use the *PHO5* promoter in *Saccharomyces cerevisiae* for which, as a unique exception, single cell data for configurations of several adjacent nucleosomes is available. This enables our detailed nucleosome dynamics modeling approach, which integrates steady state and dynamic quantities from four different data sets.

The *PHO5* promoter is a classical and extremely well-studied model system for the role of nucleosome remodeling during promoter activation (reviewed in Korber and Barbaric, 2014). It also serves as a paradigm for human promoters. The *PHO5* gene is regulated via the intracellular availability of inorganic phosphate. If phosphate is amply provided, the *PHO5* gene is lowly expressed as its promoter region is occupied by four well-positioned nucleosomes numbered N-1 to N-4 relative to the gene start. Especially nucleosomes N-1 and N-2 occlude transcription factor binding sites that are crucial for gene induction. N-1 occupies the core promoter and prevents access of the TATA-box binding protein (TBP) to the TATA-box. N-2 hinders the transactivator Pho4 from binding its cognate UASp2 element (<u>U</u>pstream <u>A</u>ctivating <u>S</u>equence <u>p</u>hosphate regulated 2), while the UASp1 element is constitutively accessible in-between N-3 and N-2. Upon phosphate depletion, a signaling cascade leads to Pho4 activation by inhibition of its phosphorylation and by increasing its nuclear localization (O’Neill et al., 1996; Komeili, 1999). Pho4 triggers an intricate nucleosome remodeling process involving up to five different nucleosome remodeling enzymes, some with redundant and some with crucial roles (Musladin et al., 2014). It also involves histone acetylation, histone chaperones and probably other cofactors that in the end lead to more or less complete removal of nucleosomes N-1 to N-5 and transcription of the *PHO5* gene (reviewed in Korber and Barbaric, 2014). Importantly, this chromatin transition was historically among the first shown to be a prerequisite and not consequence of transcription initiation (Almer and Hörz, 1986; Fascher et al., 1993; Venter et al., 1994). Thus, it strongly argued for the now widely accepted view that chromatin structure, positioned nucleosomes in particular, are not just scaffolds for packaging DNA and passive structures during genomic processes, but constitute an important level of regulation.

While the involved cofactors for this model system are known to an exceptional degree, it still remains to be understood which kind of nucleosome dynamics these cofactors bring about. To elucidate these, the full promoter nucleosome configurations in different states are needed. Indeed, for the *PHO5* promoter this information was derived from an electron microscopy (EM) single molecule scoring approach, at least for the subsystem of the N-1, N-2 and N-3 positions, in several activated, weakly activated and repressed states (Brown et al., 2013) and from single cell DNA methylation footprinting (Small et al., 2014). The N-1 to N-3 subsystem of *PHO5* promoter chromatin has been validated before to faithfully recapitulate the regulation of the full-length promoter (Fascher et al., 1993). In the single molecule *PHO5* promoter EM study (Brown et al., 2013), simple biologically motivated network Markov models were used to describe the dynamics of promoter nucleosome configurations. As the variation in nucleosome configurations is intrinsically stochastic, i.e., is not the result of other stochastic events in the nucleus (Brown and Boeger, 2014), the use of Markov models for promoter nucleosome configuration dynamics is valid. However, such models were not systematically investigated but rather several similar models that fit the data of different promoter states presented without further justification or consideration of alternative models. Other, more mechanistically detailed computational models of nucleosome remodeling with base-pair resolution using the data from (Brown et al., 2013), needed a lot of assumptions to fit the data and therefore do not represent a fully unbiased approach (Kharerin et al., 2016).

Since it is yet impossible to observe changes in nucleosome configurations at the same promoter over time *in vivo* or *in vitro*, these dynamics have to be inferred by systematic and unbiased theoretical modeling. In theory, each steady state distribution of promoter configurations can be the results of many equilibrium as well as non-equilibrium models. In equilibrium models, the free energy values of each configuration determine the steady state distribution while the reaction rates of each pair of reverting reactions are variable, as long as the ratio of rates corresponds to the Boltzmann factor of the difference in free energy. This results in zero net fluxes between different configurations in steady state, i.e. the average number of reactions in time between any two configurations is equal. In non-equilibrium models, the simple picture of an energy landscape breaks down and net fluxes in steady state are not always zero, which allows for much richer dynamics, e.g. cycles, trajectories over several configurations with same start and end, which are more likely to occur in one direction than in the other. The challenge in modeling the promoter nucleosome dynamics is to find well-motivated restrictions and assumptions to reduce the number of fit parameters to a reasonable level, while still staying unbiased, modeling on a similar level as the available experimental data and combining as many different experimental data sets as possible within the same model.

In our study, we achieved this by compiling a large and unbiased collection of possible models, mostly non-equilibrium, within our class of “regulated on-off-slide models” and selecting the models that are consistent with experimental data. Specifically, we integrate four different experimental data sets: the *PHO5* promoter nucleosome configuration data from Brown et al., 2013 of repressed, weakly activated and activated cells, data from our own restriction enzyme accessibility experiments of two different sticky (= lower accessibility) N-3 mutant promoters, to address the coupling between remodeling of the N-3 and N-2 nucleosome, and two data sets of Flag-/Myc-tagged histone exchange dynamics experiments (Dion et al., 2007; Rufiange et al., 2007) to obtain a time scale. Our regulated on-off-slide models include assembly and disassembly of N-1, N-2 and N-3 nucleosomes as well as nucleosome sliding from one occupied position to an unoccupied neighboring position. Additionally they can mimic regulated transitions from repressed over weakly activated to fully activated promoter state dynamics without changing the network topology and, once fitted, provide likely trajectories of promoter nucleosome configurations, which are completely inaccessible by experiments so far. After systematic analysis of all possible regulated on-off-slide models up to a fixed number of fit parameters, we found only very few models in agreement with all four data sets and were able to describe the effective dynamics of nucleosome configurations during chromatin remodeling at the *PHO5* promoter in yeast.

## Results

### General modeling approach

Our goal was to (i) create an unbiased class of effective minimal models featuring assembly, disassembly and sliding processes, which describe promoter configurations with the same detail as the available data sets and then (ii) investigate all models within this class to look for the least complex models to explain the data. *PHO5* promoter nucleosome dynamics can be viewed in two different levels of detail. The site-centric point of view focuses on the three positions N-1, N-2 and N-3 (in the following also referred to as N-1, N-2 and N-3 “sites for modeling purposes) and remodeling at these sites individually, i.e. without considering the nucleosome occupancy of neighboring sites (Figure 1A). An experimental example are accessibility measurements by restriction enzymes at these positions, since the measurements are taken at each position separately. A more detailed point of view keeps track of all the eight promoter nucleosome configurations (“promoter configurations), i.e. simultaneously tracks which of the three sites are occupied in each cell (Figure 1B). We used the configurational data for three mutants corresponding to different activation states of the *PHO5* promoter (“promoter states”): repressed (wild-type), weakly activated (*pho4[85-99] pho80*Δ *TATA* mutant) and activated (*pho80*Δ mutant) (Brown et al., 2013). Each promoter state shows a different steady state distribution for the occurrences of promoter configurations (Figure 1C). It is possible to calculate the three absolute site accessibilities from the promoter configuration occurrences, but important information is lost and the calculation cannot be inverted. To make full use of the information of the data of Brown et al., 2013, we based our models on the eight promoter configurations. To further restrict our models, we also used other available site-centric data, like restriction enzyme accessibility at the N-2 and N-3 site for wild-type and two sticky N-3 mutants (this study), as well as measured histone exchange dynamics at the N-1 and N-2 site (Dion et al., 2007; Rufiange et al., 2007).

**Figure 1:**
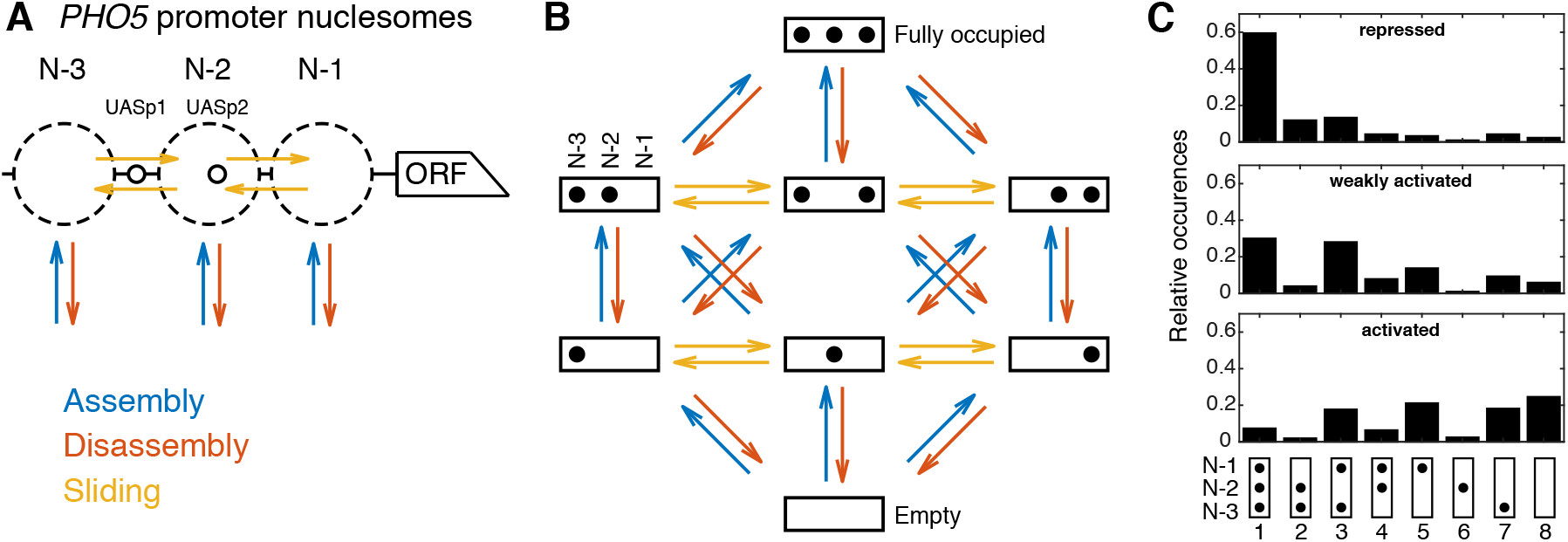
(**A**) Simplified nucleosome dynamics at the *PHO5* promoter including assembly, disassembly and sliding from a site-centric point of view. Dashed circles indicate possible nucleosome positions. Pho4 binding sites (UASp elements) are represented by small circles. (**B**) Configuration-specific modeling approach with 8 promoter configurations and 32 reactions. Arrow color code as in panel A. (**C**) Measured relative occurrences of the 8 promoter configurations indicated at the bottom as in panel B but rotated by 90° and for three different “promoter states”: the repressed wild-type, a weakly activated mutant (*pho4[85-99] pho80*Δ *TATA*) and the activated mutant (*pho80*Δ), using data from Brown et al., 2013, shown in Table S1.

### Regulated on-off-slide models

Each regulated on-off-slide model consists of a set of processes for nucleosome assembly, disassembly and sometimes sliding that allow reactions from one configuration to another (colored arrows in (Figure 1B). As each model is demanded to describe all three promoter states, at least one of these processes has to be regulated, i.e. change its rate to achieve different configuration occurrences and dynamics between different promoter states. Invoking the principle of Occam’s razor, we started with global processes and then replaced these in some cases with more specific processes, making the simplest models more and more complex until we found agreement with the considered data. A conceptually similar framework was used to model combinatorial acetylation patterns on histones, but with fixed global disassembly and without the possibilities of sliding and regulation (Blasi et al., 2016). Our approach can be applied to any system that features assembly and disassembly reactions on a fixed number of position/sites. The most important input is the steady state distribution of system configurations which can be accompanied by other known steady state or dynamic properties.

#### Global processes

The starting point was a model only consisting of two processes, global (i.e. *PHO5* promoter-wide) nucleosome assembly (“A”) and disassembly (“D”). Remodeling enzymes can bind nucleosomes and slide them along the DNA (Bartholomew, 2014; Zhou et al., 2016), so we also included optional sliding processes, which move nucleosomes to a neighboring empty site. This yields another global process, global sliding (“S”) (Figure 2A).

**Figure 2:**
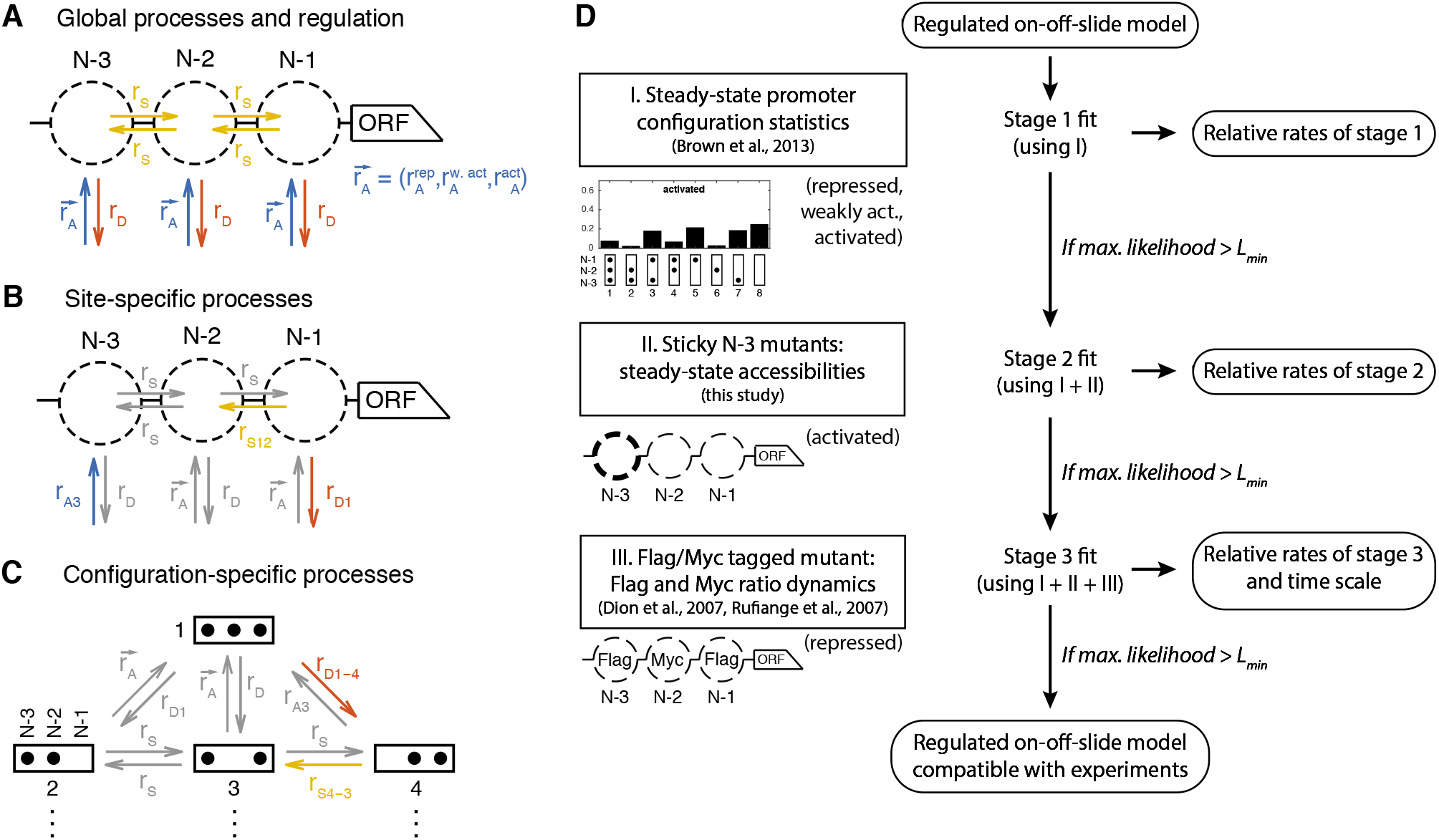
Definition and evaluation workflow of regulated on-off-slide models. On-off-slide models consist of processes from different hierarchies: (**A**) Global processes for assembly, disassembly and (optional) sliding. Global processes govern all reactions of the corresponding type with the same rate *r_A_, r_D_* or *r_S_*. To fit multiple promoter states simultaneously, some processes have to be regulated, i.e. have different rate values depending on the promoter state. In this example, the global assembly process is regulated. (**B**) Optional site-specific processes for assembly and disassembly at each position (example here with rates *r*_*A*3_ and *r*_*D*1_) and for sliding between each neighboring pair of positions (here *r*_*S*12_). Reactions in gray have not been overwritten by more specific processes (here: site-specific processes) and consequently are still determined by the rate parameters of processes on the less specific hierarchy level (here: global processes). (**C**) The last hierarchy level is given by optional configuration-specific processes governing only one reaction (here with rates *r*_*D*1–4_ and *r*_*S*4–3_). Here, only the promoter configurations 1 to 4 are shown. (**D**) Each regulated on-off-slide model is fitted and evaluated successively using the experimental data on the left-hand side (promoter states during the experiment given in parentheses). Models are discarded if they do not match the maximum likelihood threshold after each stage. With each additional experimental data set, the fit results in new optimal relative rate values of the model. Only the dynamic Flag-/Myc-tagged histone measurements enable us to also fit the time scale.

#### Regulation

We define a regulated on-off-sliding model by its set of processes combined with the information which of the processes are regulated. “Regulated” processes have more than one rate value whereas “constitutive” processes have the same rate value for each promoter state. For instance, the two simplest models have only the global assembly and global disassembly process, one being regulated and the other being constitutive. If a model is fitted to three different promoter states, one has to fit the three rate values for each regulated process, and one rate value for each constitutive process. Thus regulated processes have three fit parameters, whereas constitutive processes have one, giving four fit parameters for the two simplest models. One degree of freedom of the fit corresponds to the overall time scale of each model.

#### Site-specific assembly and disassembly processes

Brown et al., 2013 showed that only global processes are not sufficient to fit the measured occurrences of all eight configurations, even to describe only one promoter state (e.g. the activated promoter). There have to be local modifications of at least one global processes (assembly, disassembly or sliding). In Brown et al., 2013, modifications were introduced by setting certain reaction rates in the configuration network to zero, effectively searching for a good network topology in the vast discrete space of all possible topologies. In contrast, our approach has the ability to continuously deviate from the global process rate values for a given set of reactions. Here, the simplest modification is given by site-specific processes (Figure 2B). Possible biological mechanisms could be sequence-dependent effects, recruitment or inhibition of remodeling factors in a site-specific way or local differences in the epigenetic marks of nucleosomes. To incorporate such possibilities, we added site-specific assembly, disassembly and sliding processes to the pool of optional processes (see example Figure 2B). All processes, except global assembly and global disassembly, are optional and the more optional processes are allowed, the larger the number of all possible models will become. To distinguish the processes at the three nucleosome positions N-1, N-2 and N-3, we named the three site-specific assembly processes “A1”, “A2” and “A3”, and the three disassembly processes “D1”, “D2” and “D3”, respectively.

#### Site-specific sliding processes

For sliding, we included five possible processes. One process governs the rates for all sliding reactions leaving from the N-2 site to account for the possibility of start site-specific non-directional sliding (“S2*”). Two processes enable directional sliding between N-1 and N-2 sites and two directional sliding between N-2 and N-3 sites, with the short name “Sxy” for sliding from site x to site y. Note that these sliding processes actually are not only dependent on the state of one site, but two sites: the origin and the destination of the sliding process. The origin needs to be occupied and the destination to be empty. This already constitutes a correlation between neighboring sites. We still call these processes “site-specific, since they do not take into account the full configuration.

#### Modulation by more specific processes

Site-specific processes govern a subset of reactions of all reactions of a given type (assembly, disassembly or sliding) to allow deviations from the global process rate at a given position. To achieve this we invoke the following rule, which is also used for the even more specific processes of the following paragraph: If two processes govern the rates of the same reactions within a certain model, as for example the global and a site-specific assembly process, the more specific process, i.e. the one governing the rate values of less reactions, overrules the more general process, but only for these reactions. This rule allows an increased as well as a decreased rate value for the reactions of the more specific process, i.e., the specific process to be enhanced or inhibited with respect to the global process. If all reactions of a process are overruled by more specific processes in a given model, that model is redundant and will be ignored.

#### Configuration-specific processes

To allow even more specific modulations, we also added configuration-specific processes which only govern a single reaction rate and overrule any more general process for this reaction (see example Figure 2C). Since there are in total 32 reactions, this gives another 32 optional processes. For instance, the disassembly process from configuration 1 to configuration 4, is denoted with “D1-4”, sliding from configuration 4 to configuration 3 with “S4-3”.

#### Resulting model set

With the simplest regulated on-off-slide models having 4 parameters, we increased the maximal parameter number up to 7 to obtain the first models that simultaneously fit all data sets well. This yielded models with 1 regulated process having three parameter values, and 1 to 4 constitutive processes, as well as models with 2 regulated processes and 1 constitutive processes. Zero constitutive processes, i.e. only regulated processes, are not allowed, since this would correspond to independently fitting the different promoter states. After going through all combinations of processes with up to 7 parameters and ignoring effectively identical models constructed with different processes (e.g. models where one process is completely overwritten by others) we ended up with 68 145 regulated on-off-slide models in total. The relative occurrence of individual constitutive as well as regulated processes in this initial model set is the same for all assembly and disassembly processes except global assembly and disassembly and slightly lower for individual sliding processes where the number of effectively identical, and thus ignored, models is higher (Figure S1A).

#### Staging

We used the models of this new class and a step by step (“staged”) fitting procedure to determine which of them are capable to reproduce the four experimental data sets (Figure 2D). The benefit of this staging approach was that we could dissect the contributions of the different data sets to the model selection as well as the reduced model count in the later, computationally more involved, stages.

### Promoter configuration statistics

First, in stage 1, the parameter values, i.e. the rate values of the involved processes, of each model were determined by maximizing the likelihood to observe the measured *PHO5* promoter configurations (see Methods, Figure 1C and Table S1). The global assembly parameter (for the activated state) was set to 1 to fix the time scale at first, which yielded relative rates for the remaining 6 parameter values. As an example, consider the model with the processes and reaction network shown in Figure 3A. After the maximum likelihood fit, one can compare the different model nucleosome configuration distributions for the three promoter states with the configuration statistics data. Since also including the data sets discussed below into the fit did not worsen the deviations from the configuration statistics data for this specific model (data not shown), we already show here the final nucleosome configuration distributions (Figure 3B, C and D).

**Figure 3:**
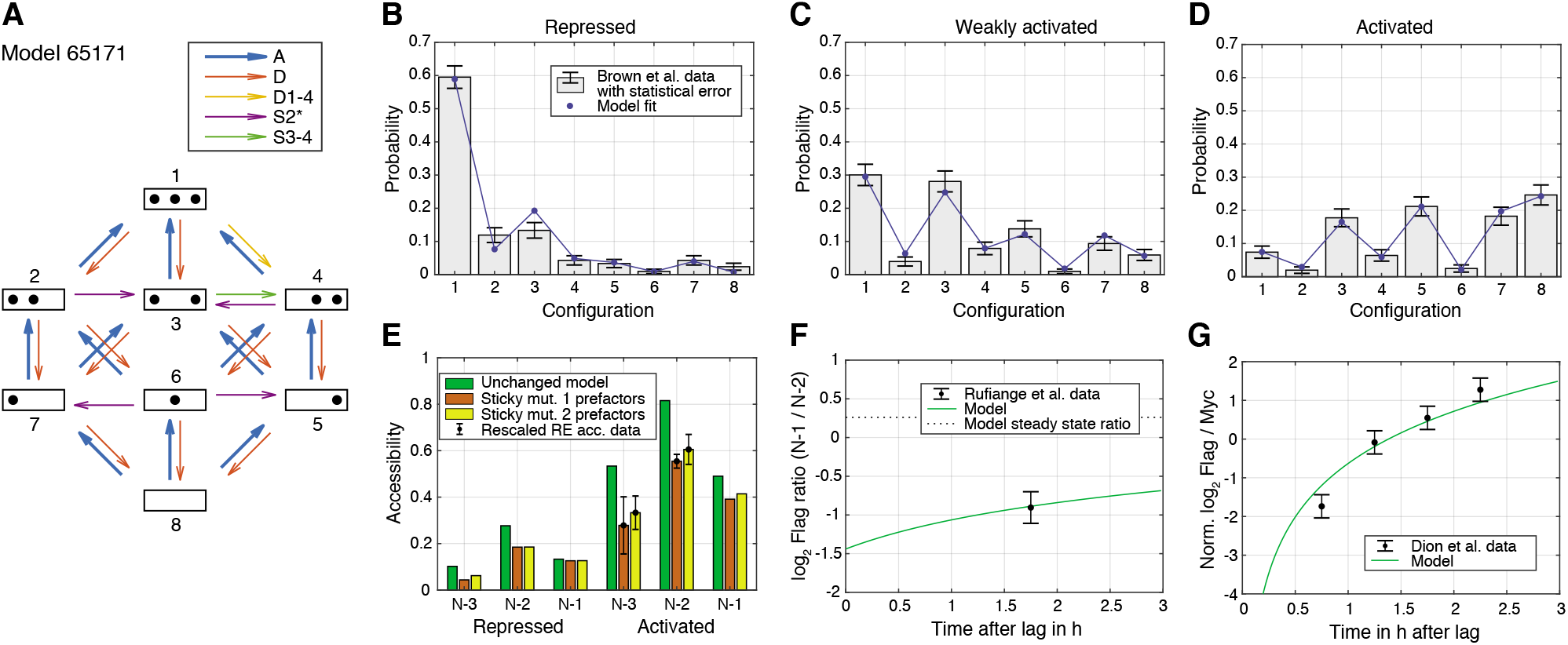
Best regulated on-off-slide model compatible with all the experimental data sets in stage 3. All fits were done simultaneously (see Methods). (**A**) Regulated on-off-slide model with the regulated process A (global assembly, thick arrows) and the constitutive processes D (global disassembly), D1-4 (disassembly from configuration 1 to 4, overrules D), S2* (sliding away from N-2) and S3-4 (sliding from configuration 3 to 4, overrules S2*). (**B, C, D**) Combined fits to the steady-state promoter nucleosome configuration occurrences in repressed, weakly activated and activated state. Only a change in the rate of the regulated global assembly process accounts for differences in the three distributions. The other processes (D, D1-4, S2* and S3-4) are constitutive and their rates do not change. The model fits in stage 1 (ignoring all other data) and stage 2 (ignoring Flag-/Myc-tagged histone exchange data) are only slightly better (data not shown). (**E**) Fit to the sticky N-3 RE accessibility data of two mutants in the activated state (error bars with standard deviation of rescaled RE accessibility). Only reaction rates involving the N-3 were allowed to divert from the previously fitted parameter values. (**F**) Fit to the Rufiange et al., 2007 data of Flag amounts at N-1 over N-2 after 2h (minus lag time) of Flag expression. The error bar corresponds to the standard deviation of two measurements. (**G**) Fit to the Dion et al., 2007 data of Flag over Myc amounts at N-1 at four time points after Flag expression. y-axis points are normalized by their mean to account for a sloppy fit parameter in the treatment of the data in Dion et al., 2007. Error bars are estimated experimental standard deviations used in the fit.

For model selection, we used the logarithmic likelihood ratio with respect to the perfect fit, *R*_1_ = −*L_I_* + *L*_0_, where *L_I_* is the maximum log10 likelihood of a given model in stage 1 and *L*_0_ the highest possible log10 likelihood to obtain the configuration data, corresponding to a perfect fit. The best model achieved *R*_1_ = 4.02 (distribution of *R*_1_ for all models in Figure S2A) and our example model in Figure 3 *R*_1_ = 4.38. We set an upper threshold to *R*_1_, *R_max_* = 6, to define models that are in good agreement with the measured configuration statistics. We used the same *R_max_* in each stage and this threshold, which denotes a likelihood ≈ 100 times lower than the current best model, gave enough room for fitting models to additional data later on. At this stage, we found 173 such models, with the top 30 models presented in Figure S3. Two of these 173 models used only 5 instead of the 6 free fit parameters (with *R*_1_ = 5.32 and 5.88, Figure S2D and Figure S4).

For comparison, Brown et al., 2013 used different network topologies to fit the different promoter states with three parameters values per state (assembly, disassembly and sliding, with assembly set to 1). The corresponding logarithmic likelihood ratio *R*_1_ of the combined model of the three states is 4.93, using effectively 6 fit parameters. This reveals the power of our systematic approach, since we found models with higher likelihood as well as models with one parameter less and not much worse likelihood.

#### Model heterogeneity

At this stage, the set of good candidate models, i.e. models with *R*_1_ below the threshold, was still quite heterogeneous, i.e., there were models where all sliding reactions have positive rates, and models with no sliding at all (Figure S2C). In fact, this was already the case for the second best model (Figure S3). Some models with *R*_1_ below the threshold did not use any configuration-specific processes (Figure S2B). The regulated processes were mostly global assembly, while some models showed regulated global disassembly or site-specific assembly at N-2 (Figure S1B). In the following stages, we further sorted out models by fitting to additional data sets.

#### Assembly-disassembly symmetry only for equilibrium models

Note that, except for a few cases, regulated on-off-slide models are non-equilibrium models. Equilibrium regulated on-off-slide models, i.e. models where the net fluxes between all promoter configurations in steady state are zero in all activation states (also known as “detailed balance” models), have an assembly-disassembly symmetry. That means, swapping all assembly processes to the equivalent disassembly process and vice versa (e.g. A to D, A1 to D1, D2 to A2), yields a model with equal maximum likelihood. Additionally, for each equilibrium model with a regulated assembly parameter there is a symmetric equilibrium model with the corresponding regulated disassembly parameter and equal maximum likelihood, and vice versa. Non-equilibrium models do not share this symmetry. Among the 68 145 investigated regulated on-off-slide models, there are 196 equilibrium models with zero net fluxes independent of their parameter values. These are models that only use global processes, where the symmetry is trivial, as well as models without any sliding nor any configuration-specific processes. These equilibrium models did not fit the data well, with the best *R*_1_ ≈ 11.

### Integration of *PHO5* promoter mutant data

#### Accessibility experiments for *PHO5* promoter mutants

In the next stages, we eliminated the regulated on-off-slide models that did not agree with further data. Small et al., 2014 published two *PHO5* promoter mutants where the DNA sequence underlying N-3 was mutated such as to increase certain dinucleotide periodicities that favor nucleosome formation and may increase intrinsic nucleosome stability (Satchwell et al., 1986). Using an at that time novel DNA methylation assay for probing nuclesome occupancies, these authors published that N-3 at these mutated *PHO5* promoters was hardly removed upon *PHO5* induction. These mutated *PHO5* promoters offered an interesting parameter modulation and we could have used, in principle, these nucleosome occupancy data for further selection among our models. However, for reasons detailed in the Discussion section, we questioned the published data and wished to check them by classical and well-documented restriction enzyme (RE) accessibility assay (Gregory et al., 1999). Therefore, it was important that we obtained the exact same strains from Small et al. and measured N-2 and N-3 occupancies at the wild type and mutant *PHO5* promoters (Table 1). We confirmed that the sticky N-3 mutant promoters show reduced removal of both N-2 and N-3 upon *PHO5* induction, but we did not confirm that these nucleosomes were hardly removed at all. Nonetheless, these data now provided additional constraints for our modeling approach as we needed to find the regulated on-off-slide models which show the same interdependence of N-2 and N-3 accessibility.

**Table 1:** Restriction enzyme (RE) accessibility of N-2 and N-3 sites in phosphate starved cells measured in this study and corresponding accessibility values of Brown et al., 2013 (RE accessibility with mean ± standard deviation of two independent biological replicates and the fold-change standard deviation calculated using standard error propagation). The sticky N-3 mutants feature manipulated DNA sequences at the N-3 site, which decrease the RE accessibility at the N-3 site compared to the wild-type. In our study, this sticky N-3 also decreases the accessibility of the N-2 site. In stage 2, we tested which regulated on-off-slide models with compatible configuration distribution in stage 1 can at the same time reproduce the accessibility fold-changes at sites N-2 and N-3 for both sticky N-3 mutants.

|  | Wild type<br>accessibility | Sticky N-3 mutant 1 |  | Sticky N-3 mutant 2 |  |
| --- | --- | --- | --- | --- | --- |
|  |  | accessibility | wt fold-change | accessibility | wt fold-change |
| N-2 RE (ClaI) | $(64.0 \pm 1.4) \%$ | $(43.5 \pm 2.1) \%$ | $0.68 \pm 0.04$ | $(47.5 \pm 4.9) \%$ | $0.74 \pm 0.08$ |
| N-2 Brown et al. | 82 % |  |  |  |  |
| N-3 RE (HhaI) | $(58.5 \pm 2.1) \%$ | $(30.5 \pm 13.4) \%$ | $0.52 \pm 0.23$ | $(36.5 \pm 7.8) \%$ | $0.62 \pm 0.13$ |
| N-3 Brown et al. | 55 % |  |  |  |  |

#### Accessibility fold-changes

The Small et al., 2014 strains included corresponding isogenic wild type strains and we noted that our RE accessibilities for wild type in the activated state were lower than the accessibilities in the activated state of the wild type calculated from the data from Brown et al., 2013 (Table 1). This discrepancy likely stems from different experimental conditions. The Brown et al., 2013 study used a different strain background, YS18, and the *pho80* allele in high phosphate conditions for induction, while we used the S288C background and over night phosphate starvation to achieve direct comparison with the data by Small et al., 2014. We saw repeatedly that S288C strains do not yield as high ClaI accessibility values in the induced state as the YS18 background, which was formerly used in classical *PHO5* studies reporting such high degree of *PHO5* promoter nucleosome remodeling (Ertel et al., 2010 and data not shown). Nonetheless, this discrepancy did not matter for our purposes as we could use the accessibility fold-changes of mutants compared to the wild-type to test our models in order to normalize the accessibility values coming from different experiments.

#### Modeling approach

In stage 2, we fitted the experimental fold-changes together with the data used in stage 1, minimizing *R*_2_ = − (*L_I_* + *L_II_*) + *L*_0_ with *L_II_* being the log likelihood to obtain the accessibility fold changes (see Methods). We needed to systematically test for each model whether reasonably large changes in reaction rates involving the N-3 nucleosome could lead to the same behavior seen in the experiment. Since we did not know the exact consequences of the sticky N-3 mutation on the reaction rates or how to translate them within a given model, we had to consider all possibilities of changes in reactions where the N-3 nucleosome is involved. We achieved this by including prefactors for these 12 reaction rates (Figure S5), which were fitted together with the model parameters to the configuration statistics data and the accessibility fold changes of both sticky N-3 mutants and allowed each prefactor to vary between 1/5 and 5 for each sticky N-3 mutant. If we interpreted the ratio of a pair of reverting reaction rates by *e*^Δ*E_f_*/k_B_T^, this would lead to a maximal possible absolute change in nucleosome binding energy of Δ*E_f_* ≈ 3.2 k_B_T ≈ 1.9kcal/mol (ignoring all other reactions and if both reactions have non-zero rates). We decided to use 4 prefactors per sticky N-3 mutant, one for each group of reactions, assembly at N-3, disassembly at N-3, sliding from N-3 to N-2 and from N-2 to N-3 (Figure S5), respectively.

Using the same likelihood threshold as in stage 1 left us with 15 models that fit both sticky N-3 mutants well (Figure S6A), with the best logarithmic likelihood ratio of *R*_2_ = 4.38 for the model in Figure 3, with Figure 3E showing a perfect fit to the sticky N-3 fold changes (also including the following data sets). Out of these 15 models, only three models without configuration-specific processes remained (Figure S6B). With this fit we also excluded models without sliding processes (Figure S6C) as well as models with less than seven fit parameters (Figure S6D).

#### Prefactor behavior

The fitted prefactor values show stable qualitative behavior for the models with *R*_2_ below *R_max_* in both mutants (Figure S7): consistent with the reduced experimental accessibility at N-3, assembly at N-3 is increased by a prefactor greater than 1 while disassembly at N-3 is decreased by a prefactor smaller than 1. Furthermore, in these models the sliding from N-3 to N-2 is increased and the sliding from N-2 to N-3 decreased by the two sliding prefactors. While these favor the accessibility at N-3 again, they lead to the concomitant accessibility decrease at N-2.

#### Relative occurrence of sliding

Taking into account the sticky N-3 data has a strong impact on the relative occurrences of sliding processes, as it strongly favors models with either S32 or S3-4 processes, both of which enable the sliding from N-3 to N-2 (Figure S1C). All 15 good models in stage 2 include at least one sliding process (Figure S8).

### Flag-/Myc-tagged nucleosome exchange simulation

In stage 3, we used histone dynamics measurements to further constrain our models and, for the first time, were able to determine the optimal time scale of each model. Histone dynamics measurements reflect the appearance/disappearance of nucleosomes regardless if via assembly/disassembly or sliding and is measured in cells that constitutively express Myc-tagged histones and are then induced to express also Flag-tagged histones, which, over time, are incorporated into nucleosomes (Schermer et al., 2005). We used the histone pool model as well as the measured average Flag over Myc amount ratio at the N-1 in MNase-ChIP-chip assays of Dion et al., 2007 and the measured Flag at N-1 over Flag at N-2 amount ratio in Flag-tagged H3 MNase-ChIP-qPCR assays of Rufiange et al., 2007. We investigated the nucleosome configuration dynamics of each model by keeping track which nucleosome contains a Flag- or Myc-tagged histone H3 (see Methods). This enabled us to calculate the dynamics of the ratio of Flag at N-1 over Flag at N-2 amount (Figure 3F and Figure S9) and the ratio of Flag over Myc amount at the N-1 (Figure 3G and Figure S10) for each model and determine the agreement with the two histone dynamics data sets. We obtained 7 models in agreement with the data, with a best logarithmic likelihood ratio of *R*_3_ = − (*L_I_* + *L_II_* + *L_III_*) + *L*_0_ = 4.61, with *L_III_* being the summed log10 likelihoods of the third and fourth data set. Thus, the threshold value of *R_max_* = 6 denotes a likelihood ≈ 25 times lower as the best achieved value of *R*_3_.

### Properties of satisfactory models

#### Processes and rate values

Out of the 68 145 models with at most 7 fitted parameter values, 7 models satisfied our threshold for the likelihood in the combined fit to the nucleosome configuration data, the sticky N-3 mutant accessibility data and the H3 exchange data. These “satisfactory” models, their processes and rate values are presented in Figure 4, with the corresponding reaction rates shown in Figure S11.

**Figure 4:**
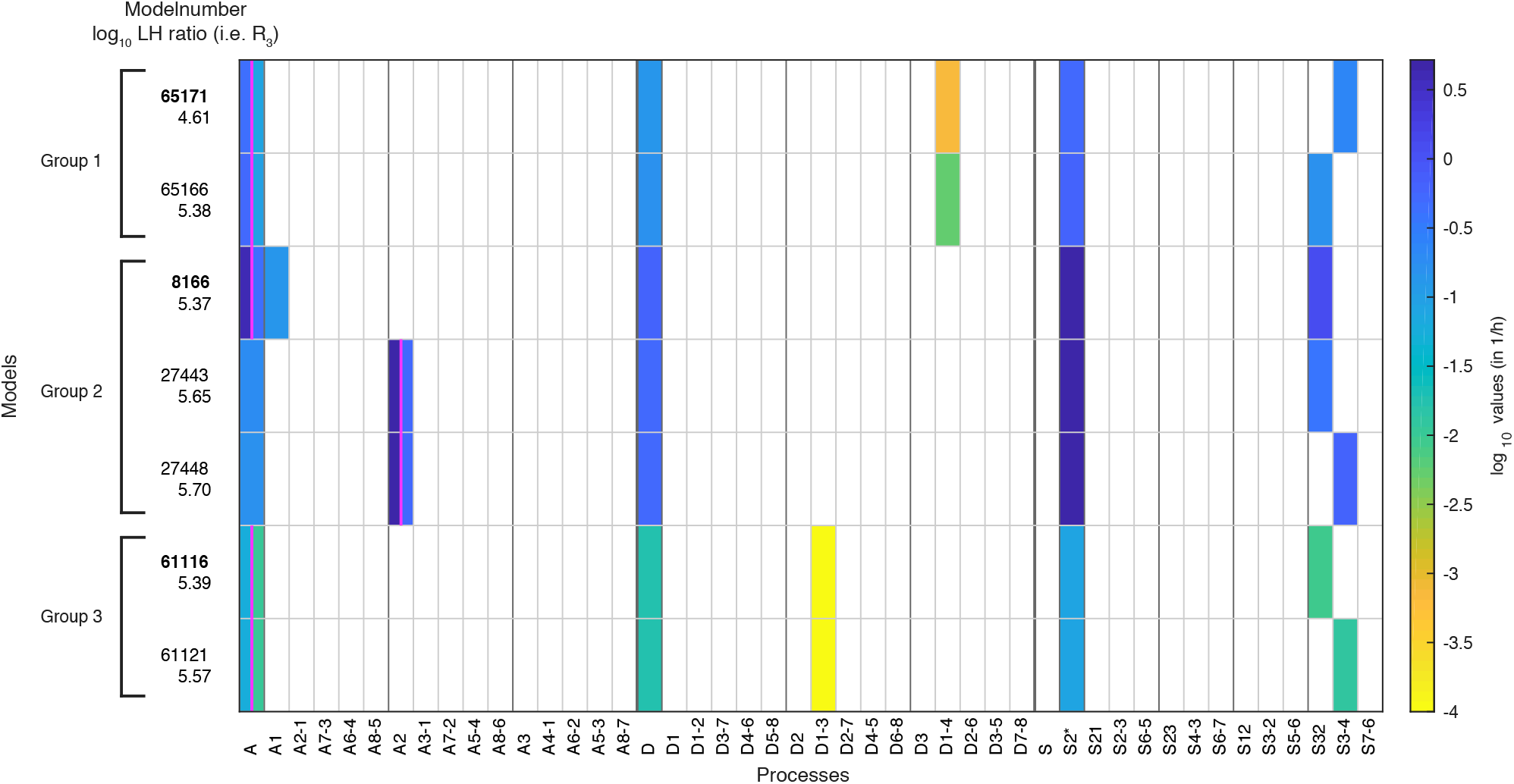
The 7 models that agree with all four data sets after stage 3. The colored boxes in each row show the model processes and their rate values. White boxes denote the absence of a process in a model. Regulated processes are separated into two differently colored boxes for repressed (left half) and activated (right half) promoter state. Weakly activated rate values are not shown here. The exact rate values are printed in Table S2 and Table S3. On the left side are the model number and the log10 ratio of the best possible likelihood and the model likelihood, *R*_3_. The models are grouped with respect to similarities in the site-centric net fluxes (Figure S16). The groups’ representatives are printed in bold with net fluxes compared in Figure 5.

#### Only one regulated process

All satisfactory models had one regulated process and 4 constitutive processes (Figure 4). Simultaneous regulation of two processes in combination with 1 constitutive process (also summing up to 7 fit parameters) did not fit the data well enough, probably because regulation with 2 processes leaves only 1 possible constitutive process slot and only three processes in total. Thus, the rather simple regulation of only one process is preferred over regulation of several processes, when keeping the total number of fit parameter constantly at 7.

#### Regulation by assembly

Within the 7 satisfactory models, the regulated processes are global assembly, A (5x) and assembly at N-2, A2 (2x) (Figure 4 and Figure S1D). The fact that regulation by assembly rather than disassembly is more likely to reproduce the data can already be observed after the maximum likelihood fit to the configuration data in stage 1 (Figure S1B). The rates of all regulated assembly processes decrease when going from the repressed over the weakly activated to the activated state. This is in agreement with the total nucleosome occupancy decrease on the promoter.

#### Sliding processes needed

Since all sliding processes are optional, we asked whether sliding is needed to fit all data sets. We found that sliding occurs in all 7 models, and in each case with at least two different processes, i.e. just global sliding with the same rate for each sliding reaction is not sufficient. Every satisfactory model employs the sliding process away from N-2 (S2*) and a process that allows sliding from N-3 to N-2 (Figure 4 and Figure S1D).

#### Use of site- and configuration-specific processes

On-off-slide models combine processes from different hierarchies. Each model has a global assembly and global disassembly process, but global sliding is optional and not used in any of the satisfactory models. Each of these models has one to three site-specific processes out of A1, A2, S2* and S32, which overwrite global processes. We found two models without configuration-specific processes, models 8166 and 27443 (Figure 4). All other satisfactory models have one or two configuration-specific processes. Thus, configuration-specific processes are helpful, but not needed to achieve agreement with the data. However, that does not mean that the three nucleosome sites can be independent of each other, since the sliding reactions always introduce coupling between neighboring sites.

#### Fluxes

Knowing all reaction rates, it is easy to calculate the directional fluxes (Figure S12 and Figure S13) as well as the net fluxes (Figure S14 and Figure S15) between all configurations for each satisfactory model and for different promoter states. Since there are no methods available to measure these fluxes experimentally, modeling approaches provide the only view into possible internal promoter configuration dynamics. As stated before, the vast majority of all regulated on-off-slide models are non-equilibrium models, i.e. have non-zero net fluxes. In the repressed promoter, the seven satisfactory models showed the highest fluxes as well as net fluxes occurring between the configurations with three or two nucleosomes (Figure S12 and Figure S14) and cyclic net fluxes from configuration 1 to 2 to 3 and back to 1. Despite qualitative similarities, the fluxes and net fluxes also show the different behavior of the seven models, for example only five of them showed cyclic net fluxes from configuration 1 to 4 to 2 and back to 1. In the activated promoter state, the higher fluxes and net fluxes between the first four configurations are lost and the differences between the seven models become more pronounced.

#### Site-centric net fluxes

Going back to the view point of nucleosome sites as in Figure 1A, we define effective “site-centric net fluxes” by summing all assembly/disassembly net fluxes at each site and sliding net fluxes between N-1 and N-2 as well as N-2 and N-3 (Figure S16). The site-centric net fluxes give a simplified picture of the net paths of the nucleosomes on the promoter, but ignore the events on neighboring sites, which are only correctly depicted in the directional or net fluxes between the promoter configurations. While in all satisfactory models the N-2 site had a central role with the strongest net nucleosome influx in all promoter states, we also found differences and divided the satisfactory models into three groups. 2 models (group 1) have site-centric net influx only at N-2 and N-3 for the repressed state and only at N-2 for the activated state. The other models have site-centric net influx only at N-2 for all promoter states, but show very different flux amounts. 3 models (group 2) show a repressed N-2 site-centric net influx of ≈ 0.59 h^−1^, while the last 2 models (group 3) have ≈ 0.013 h^−1^. The time scale parameters of group 3 are 10-fold lower than for the other two groups and have larger error bars (see Methods and Table S2), allowing the time scale to increase by up to a factor of 10 as well as becoming arbitrarily small, while still meeting the fit threshold. Thus, the time scale parameter of this group is not properly determined by the given data. For each group, we picked a representative model (highest likelihood within the group) and show the net fluxes and the site-centric net fluxes in Figure 5.

**Figure 5:**
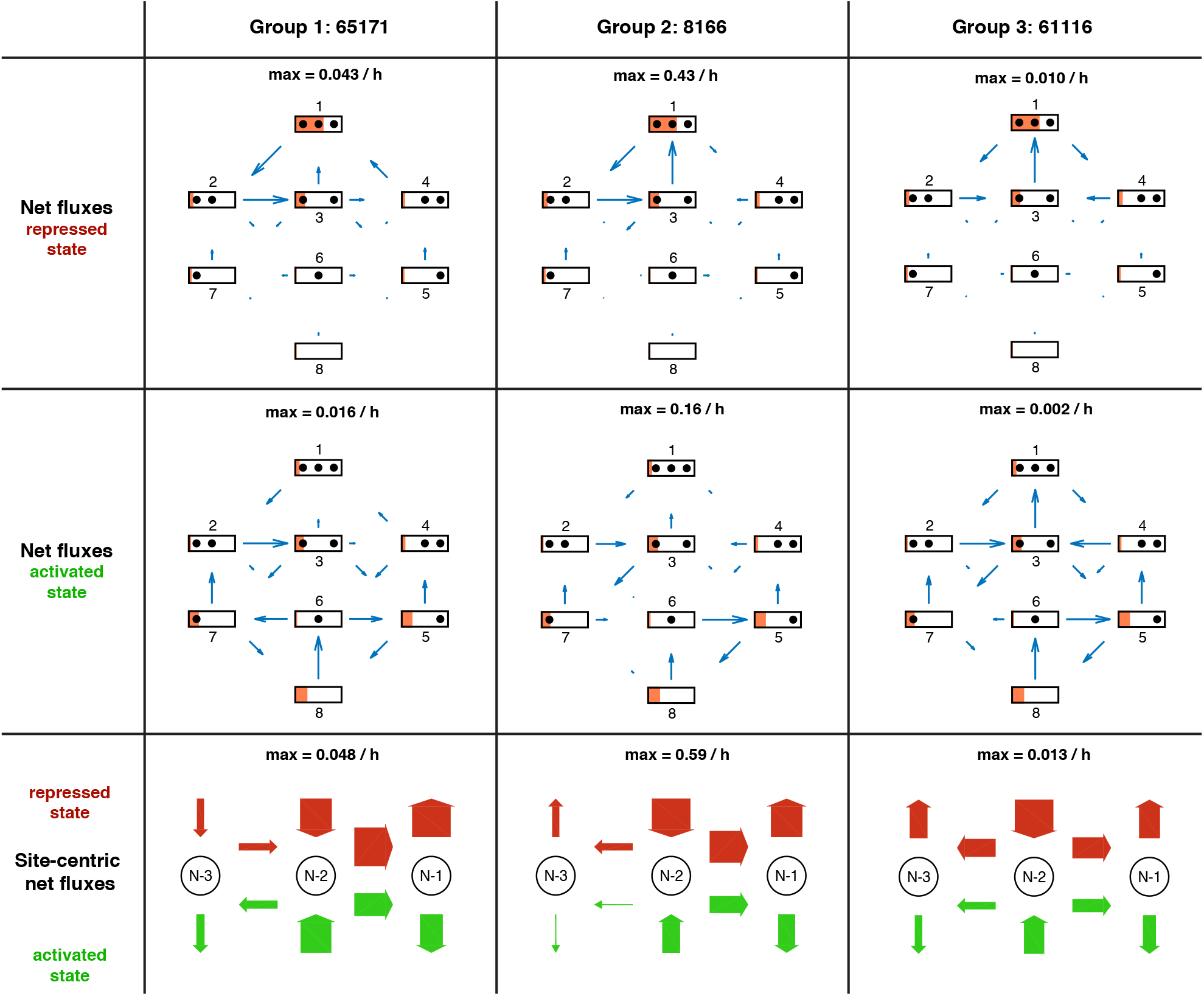
Overview of net fluxes and site-centric net fluxes for the three group representatives. First two rows: net fluxes in repressed and activated promoter state (arrows) with configuration probabilities as orange horizontal bars (a filled promoter rectangle corresponds to probability 1). Arrow length indicates the relative flux amount within a flux network with the maximum stated above. Third row: site-centric net fluxes in repressed (red) and activated (green) state, obtained by summing all assembly/disassembly net fluxes at each site and sliding net fluxes between N-1 and N-2 as well as N-2 and N-3. Here the arrow thickness indicates the amount of flux with the maximum stated above.

#### Maximal reaction rates

After setting the time scale for each model, we investigated the rate values for each reaction (Table S2 and Table S3). Within all satisfactory models, the highest rate of assembly and disassembly processes were 5h ^−1^ and 0.6 h^−1^, respectively. Sliding rate values had a maximum of 5h^−1^, corresponding to 0.2bp/s assuming an unidirectional travel of 160 bp with constant speed between two sites. These speeds are within the capabilities of yeast ySWI/SNF or RSC remodeling complexes with translocation speeds of up to 13 bp/s (Zhang et al., 2006).

#### Bounds for chromatin opening and closing times

*PHO5* induction results from consecutive signal transduction, promoter chromatin opening, transcription initiation and downstream processes that lead to functional Pho5 acid phosphatase gene product. On the level of *PHO5* mRNA or acid phosphatase activity, induction starts about two hours after phosphate starvation of the cells (Rajkumar et al., 2013; Barbaric et al., 2001), while after the same time chromatin opening in terms of chromatin accessibility is usually complete, i.e. the kinetics of *PHO5* promoter chromatin opening are faster (Schmid et al., 1992; Barbaric et al., 2001; Korber et al., 2006; Barbaric et al., 2007). We modeled here *PHO5* promoter chromatin remodeling and can give approximate upper bounds for the chromatin opening rates for each regulated on-off-slide model (“effective chromatin opening rate”, see Methods).

Models in group 1 had an effective chromatin opening rate close to 0.2 h^−1^. Group 2 showed a faster effective chromatin opening rate of ≈ 0.8h^−1^, while group 3 yielded ≈ 0.02h^−1^. Since these values are proportional to the time scale, they inherit its uncertainty. The effective chromatin opening rate of group 2 was high enough to reflect the experimentally measured kinetics. The other two groups could also match these kinetics, although just barely, if the maximum time scale uncertainty was considered (approximately an additional factor of 2 for group 1 and a factor of 10 for group 3, see Table S2).

Conversely, we did the same calculations for the “effective chromatin closing rate”, which is an upper bound of how fast a given model can switch to the repressed state. The ratio of the effective chromatin closing rate over the opening rate was between 2.9 and 3.5 for all satisfactory models. Even just after the first fit in stage 1, all investigated assembly-regulated models in agreement with the configurational data had a ratio ranging from 1.5 to 4.5. The few disassembly-regulated models after stage 1 had a ratio ranging from 0.25 to 0.65. This further supports the assembly-regulated models as promoter chromatin closing and repression of *PHO5* transcription were experimentally shown to be faster than chromatin opening and transcription activation. After the shift from phosphate-free to phosphate-containing medium, repression of *PHO5* transcription was almost complete within 20min (Schermer et al., 2005) and 65% of chromatin closing was achieved within 45 min (Schmid et al., 1992).

#### Extending the threshold

The properties of satisfactory models remained stable upon increasing the logarithmic likelihood ratio threshold *R_max_* from 6 to 7. This yielded 28 models with *R_3_ < *R*_max_* (data not shown). A notable exception was the appearance of models with both, global sliding and sliding away from N-2 (S2*) and the first model with only one sliding process (S2*). Otherwise, they all showed very similar properties as the models selected with *R_max_* = 6.

## Discussion

We present a new approach for modeling nucleosome dynamics at promoters: regulated on-off-slide models. They include nucleosome assembly, disassembly and sliding and enable a combined fit of data for different promoter activation states. The hierarchical approach of global, site-specific and configuration-specific processes allowed us to represent different network topologies, i.e., switching “off” reactions by using a very low rate of a non-global process, and continuous variations among network topologies. We investigated all regulated on-off-slide models with up to 7 fit parameters and achieved a completely unbiased modeling approach that provides insight into nucleosome dynamics, which are so far not accessible by experiments.

Using the *PHO5* promoter as example, regulated on-off-slide models provided a unique integration of four different data sets in nucleosome resolution: multi-nucleosome configuration measurements, our own nucleosome accessibility experiments in sticky N-3 mutants and two Flag/Myc-tagged histone H3 exchange experiments.

While we obtained and used the same two sticky N-3 mutant strains as published (Small et al., 2014) we did not use their published chromatin data. These were generated by an at that time novel single molecule DNA methylase footprinting approach and seemed less trustworthy for the following reasons. First, it was not clear if the DNA methylation reactions were saturated (for a detailed discussion see Oberbeckmann et al., 2019). Second, accessibility at the N-3 position in wild type cells was lower (7%) in the activated compared to the repressed (28%) state, which is in contrast to chromatin opening upon activation and the data by Brown et al., 2013 with 55% accessibility at N-3 in the activated promoter state. Third, the authors did not detect any case where all three N-1 to N-3 nucleosomes were removed upon activation, although this configuration was among the most populated in the study by Brown et al., 2013. Both the second and third point may be due to incomplete methylation and/or to erroneous interpretation of methylation footprint patterns as not only nucleosomes but also other factors like the transcription initiation machinery could obstruct methylation. Fourth, we found it hard to believe that the point mutations introduced in the sticky N-3 *PHO5* promoter mutant versions would completely abolish chromatin opening not only at the N-3 but also at the N-2 nucleosome as claimed by the authors (Small et al., 2014).

Therefore, we rather employed the classical and well-documented restriction enzyme accessibility assay (Almer et al., 1986; Gregory et al., 1999) to monitor *PHO5* chromatin opening in the sticky N-3 mutants. In this assay, both sticky N-3 mutants still showed considerable chromatin opening but confirmed the general conclusion by Small et al., 2014 that opening of both the N-2 and N-3 nucleosomes was less extensive than for the wild type promoter. Even though the effect was less pronounced than claimed originally, it is remarkable how it substantiates earlier observations at the yeast *PHO8* and *PHO84* promoters (Wippo et al., 2009), which are co-regulated with the *PHO5* promoter, about the role of underlying DNA sequence in stabilizing nucleosomes against remodeling. Recent *in vitro* experiments started to reveal how chromatin remodeling enzymes are regulated by nucleosomal DNA sequences (Lorch et al., 2014; Winger and Bowman, 2017; Krietenstein et al., 2016). The *PHO5* promoter periodicity mutants pioneered by Small et al., 2014 may provide an impressive case how this is relevant *in vivo*.

Keeping these four data sets in mind, we designed our models to consider full nucleosomes, not individual histones, during assembly and disassembly and used an effective description over a more detailed approach with base-pair resolution and transcription factor dynamics (Kharerin et al., 2016). In this way, steady state data combined with dynamical data provided new insights into the nucleosome configuration dynamics. We arranged our regulated on-off-slide models such that at least one process was regulated, i.e. its rate value varied depending on the promoter state. This gave us the opportunity to examine how regulation between different promoter states was most likely achieved. We found that regulation from repressed over weakly activated to activated promoter states can be surprisingly simple. Only using the configurational data of Brown et al., 2013 was not enough to restrict our model set. For instance, the best and second best models after stage 1 had almost identical likelihood to reproduce the data, but very different properties, as the first was regulated by global assembly and used a sliding process while the second was regulated by global disassembly and had no sliding at all.

Out of 68 145 tested models only 7 models satisfied our threshold criterion after fitting all four data sets. All 7 satisfactory models enabled sliding away from the N-2 position, but towards the N-2 only from the N-3 position. Equilibrium models and models without any sliding within the tested class did not fit all four data sets. As we made all sliding processes optional, this showed that sliding is essential and has a net directionality. Configuration-specific processes, i.e. processes governing only a single reaction, were not needed in all but two of the 7 models, and could increase the likelihood. This also means that in these two cases, the coupling between positions introduced by sliding was sufficient to reproduce the experiments.

By incorporating dynamical Flag-/Myc-tagged histone exchange data sets we were able to set the time scale for each model, which can not be done by using steady state data only. This is a completely new use case for these data sets and we took care not to over-interpret them. We found that some models had a rather “sloppy” time scale, i.e. the time scale could vary strongly without notably decreasing the fit quality. One reason was that the ratios at different times of Flag over Myc amount at N-1 needed to be shifted by their mean to take into account the high fit error of the absolute values in Dion et al., 2007. The method of Rufiange et al., 2007 did not have this problem, but unfortunately gave us measurements at only one time point, not restricting the slope of the dynamics of the ratio of N-1 and N-2 Flag amounts Figure S9.

Our models were selected according to four independent data sets derived from three orthogonal experimental approaches. Importantly, given the time scales, we could estimate lower boundaries on how quickly the satisfactory models could switch from a closed state to an open state and vice versa. Taking into account the time scale errors, the effective chromatin opening rates were compatible with experimental values and the comparison between effective chromatin opening versus closing rates were only compatible with the regulated assembly rather than disassembly models already after the first stage just using the data from Brown et al., 2013. This amounts to confirmation by a fourth type of orthogonal data.

While the identification of essential and directional sliding during *PHO5* promoter chromatin opening, the estimated time scales and the central role for the N-2 nucleosome in all seven models matched expectations based on earlier studies, we were surprised that only a global or site-specific (N-2) regulated assembly, but not a regulated disassembly process was compatible with the data which would be commonly expected. In case of an equilibrium system, the same chromatin opening could result from more disassembly or from less assembly. However, as mentioned above, our selected models are non-equilibrium models and the tested equilibrium models did not fit the data at all. Therefore, the selection of regulated assembly does not just represent the flip side of the common view but seems indeed surprising. We note that models with regulated disassembly could result in satisfactory fits to all data sets after we allowed one additional fit parameter, but were still a very small minority among all satisfactory models, with best maximum likelihood drastically lower (less than 1/20) than the best assembly regulated models. However, with more parameters the modeling approach becomes less distinctive and a large number of different mechanisms could be modeled and fitted successfully. Therefore, at first sight, regulated assembly counters the common view on promoter chromatin opening mechanisms as derived from pioneering studies at the yeast *PHO5* (Korber and Barbaric, 2014) or other, like the *HO* (Cosma et al., 1999) promoter, but also at mammalian promoters like the glucocorticoid-regulated MMTV promoter (Deroo and Archer, 2001; Johnson et al., 2008). According to this view, chromatin opening is triggered by binding of a (transcription) factor that locally recruits, either directly or via histone modifications like acetylation (Hassan et al., 2001), a chromatin remodeler, which in turn mediates nucleosome removal either by sliding and/or by disassembly (Sudarsanam and Winston, 2000; Vignali et al., 2000; Workman, 2006; Becker and Hörz, 2002; Narlikar et al., 2002). It is mainly based on i) experiments showing physical interactions between transcription factors and remodelers (Neely et al., 1999; Yudkovsky et al., 1999; Natarajan et al., 1999) or between remodelers and modified chromatin (Hassan et al., 2001), ii) on chromatin immunoprecipitation or microscopy data showing transcription factor or histone modifier recruitment to the promoter upon promoter activation (Cosma et al., 1999; Kowenz-Leutz and Leutz, 1999; Barbaric et al., 2003; Dhasarathy and Kladde, 2005; Johnson et al., 2008), and iii) on *in vitro* assays demonstrating nucleosome sliding and disassembly activities for chromatin remodelers (Tsukiyama et al., 1994; Lorch et al., 1999; Längst et al., 1999; Hamiche et al., 1999). As chromatin opening amounts to the net removal of nucleosomes it was always intuitive to see nucleosome disassembly as the regulated process. Nonetheless, we note that remodelers were equally shown to mediate nucleosome assembly *in vitro* (Lorch et al., 1999; Haushalter and Kadonaga, 2003) and wonder if the well-documented remodeler recruitment at promoters upon promoter activation may also result in downregulation of nucleosome assembly rather than upregulation of disassembly.

We suggest to reconsider the common view for the case of the *PHO5* promoter in the light of our results here and in the context of a long standing and recently revived debate regarding the role of binding competition in nucleosome remodeling. Before ATP dependent remodeling enzymes and their active nucleosome displacement activities were fully recognized, it was proposed that binding competition between sequence specific DNA binders like transcription factors and the histone octamer, i.e. mass action principles, was a major mechanism for nucleosome removal. For example, if a nucleosome was positioned over Gal4 binding sites in the absence of Gal4 it could be displaced by adding Gal4 *in vitro* (Workman and Kingston, 1992) or inducing Gal4 *in vivo* (Morse, 1993). Recently, the class of pioneer factors and general regulatory factors that have important roles in opening chromatin or keeping chromatin open, e.g., in the context of cellular reprogramming, were also suggested to displace nucleosomes by binding competition (Yan et al., 2018; Donovan et al., 2019; Iwafuchi et al., 2020). For the *PHO5* promoter, this mechanism seemed attractive as its transactivator Pho4 indeed competes with nucleosome N-2 during binding to the intranucleosomal UASp2. Further, a *PHO5* promoter lacking UASp2, i.e., left only with the constitutively accessible UASp1 in-between N-2 and N-3, was not induced during phosphate starvation *in vivo* (Fascher et al., 1990). However, early studies dismissed an essential role for binding competition at the *PHO5* promoter as i) the coregulated *PHO8* promoter was opened without an intranucleosomal UASp (Barbarić et al., 1992), ii) even the ΔUASp2 *PHO5* promoter mutant could be opened if *PHO4* was overexpressed (Fascher et al., 1990) and iii) even an overexpressed Pho4 version containing a functional DNA-binding but no transactivation domain could not displace N-2 and trigger chromatin opening although just the Gal4 DNA binding domain could displace nucleosomes in other contexts (Workman and Kingston, 1992; Morse, 1993). However again, binding competition at UASp2 in N-2, while not essential if binding to UASp1 was boosted by increasing its affinity or by *PHO4* overexpression, did have a critical role in *PHO5* promoter chromatin remodeling for wild type UASp1 and *PHO4* expression levels (Ertel et al., 2010). Therefore, we suggest for the wild type version of the *PHO5* promoter with regard to Pho4 binding to UASp elements, which we exclusively studied here, that the competition between Pho4 binding to the promoter and nucleosome assembly at the promoter corresponds to the downregulated assembly process upon promoter activation as revealed by our modeling approach. Impeding the assembly of N-2 by Pho4 binding at UASp2 could directly correspond to the downregulated N-2 assembly processes in our models 27443 and 27448 (Figure 4). Pho4 binding to both UASp2 and UASp1 may correspond to impaired global assembly in the other models. Such a promoter chromatin opening mechanism via blocking nucleosome assembly through binding competition does not preclude that Pho4 recruits chromatin remodelers that are important for the nucleosome dynamics during the chromatin transition, as posed by the common view, but it shifts the interpretation of previous data away from a focus on a regulated disassembly to a regulated assembly process. Therefore, our models are not contradicted by any existing evidence, but may prompt us to reconsider the principal direction of nucleosome dynamics *in vivo*, which so far could not be addressed by any approach. In hindsight, even though *PHO5* promoter opening by *PHO4* overexpression in the absence of UASp2 led to the same open promoter state as judged by DNAseI indirect endlabeling and restriction enzyme accessibilities (Fascher et al., 1990), different effective nucleosome dynamics may underlie this mutant compared to the wild type promoter transition. If the same kind of single molecule nucleosome configuration data as in Brown et al., 2013 were available for this ΔUASp2 *PHO5* promoter chromatin transition, our modeling approach could test this, especially if a regulated assembly process was coupled to binding competition at N-2. It was maybe not justified to assume that the *PHO5* promoter chromatin opening mechanism was the same for different promoter mutants just because the final result was the same. This may exemplify even the more how blind we were to the underlying effective nucleosome dynamics by just monitoring start and end states of a chromatin transition.

## Materials and Methods

### Maximum likelihood fits

Depending on the stage of the analysis, we optimized the parameter values, i.e. the rate values of the processes within a given model, 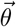, by maximizing the sum of log10 likelihood values: 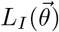 in stage 1, 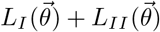 in stage 2 and 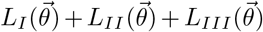 in stage 3. Note that the optimal 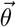 can differ between different stages. We ignored additive constants to the log likelihood, so that after including the next data set into the fit, a perfect agreement between model and the additional data set, already with the 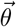 from the previous stage, would lead to the same log likelihood value as before. In all stages we used Matlab’s fmincon function to find the parameter values 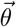 that maximize the likelihood. In the following, 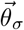 denotes the vector of all constitutive parameter values and the regulated parameter values of promoter state *σ*. Thus the entries of 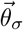 are a subset of the entries of 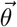.

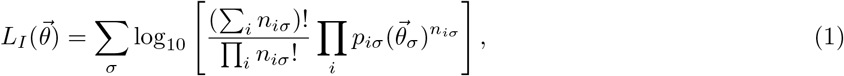

with *n^iσ^* being the number of observations (data from Brown et al., 2013), and 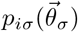 the steady state probability of the model with parameter values 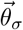, of promoter configuration *i* in promoter state *σ*. I.e. 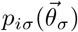 is the solution of 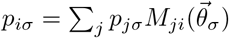.

Here 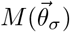 is the transition rate matrix for promoter state *σ*. A non-diagonal entry 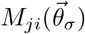 is the rate to go from configuration *j* to configuration *i* and is non-zero only for valid assembly, disassembly and sliding reactions and then given by the entry of 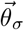 which holds the parameter value of the process that governs this reaction in the given model. Diagonal entries are given by 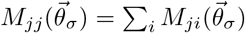.

For each model we used 100 different sets of initial parameter values to ensure we have found a robust maximum. To calculate the steady state distribution of a given model for fixed parameter values we used the state reduction algorithm (Sheskin, 1985; Grassmann et al., 1985; Shanbhag and Rao, 2003). We limited the range of parameter values to [10^−2^; 10^2^], with 1 being the rate value of the global assembly process for the activated state. A wider range of [10^−3^; 10^3^] did not affect the results. In 6.46% of all models the 100 tries found at least two different maximal likelihood values, which were always extremely likelihood values. In 3.72% of all models, the found maximum likelihood parameter values were not unique. In both cases, none of these problematic models are among models with relatively high maximal likelihoods.

From stage 2 on we also used

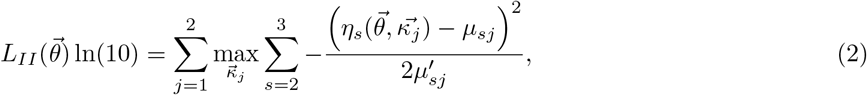

with *μ_sj_* and 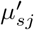 being mean and variance, respectively, of the measured accessibility fold changes in active state of mutant *j* at nucleosome site *s* (2 for N-2 and 3 for N-3), 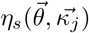 being the corresponding model fold change, and 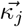 being the four values of the rate prefactors of sticky mutant *j*. Strictly speaking *L_II_* depends only on 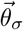, with *σ* being the activated state. Note that the exact values of prefactors found during optimization depended on their initial condition, as their best values were often sloppy or even not unique, but still resulted in the same minimal fit error.

For stage 3, *L_III_* has two contributions, one for each histone H3 exchange experiment:

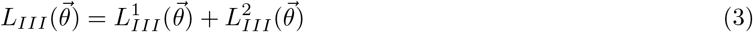

For the first contribution, to fit the data from Rufiange et al., 2007, we used

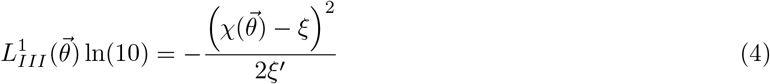

with *ξ* and *ξ*′ being the mean and variance, respectively, of the measured log2 ratios of Flag amounts at N-1 over N-2 (Rufiange et al., 2007, ratio values 0.591 and 0.483 for replicate 1 and 2, respectively) and 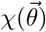 the corresponding log2 ratio of the model (see Method Section H3 histone exchange model). Strictly speaking *L_III_* depends only on 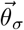, with *σ* being the repressed state.

For the second contribution, to fit the data from Dion et al., 2007, let *ν_j_* denote the log2 ratios of Flag amount over Myc amount at N-1, with *j* = 1, 2, 3, 4 indicating the four different time points. We obtained these from fitting the normalization constant of each time point using the raw data (mean values of two replicates) as described (supplementary material in Dion et al., 2007, with the nucleosome pool parameters as in the section below) and then used the results of the probe at the N-1 position of the *PHO5* promoter. Unfortunately, neighboring probes were only in linker regions between promoter nucleosome positions. As mentioned in Dion et al., 2007, the absolute values of *ν_j_* have large errors (due to a sloppy global normalization constant), while the differences between time points were determined with reasonable accuracy. As a reference we chose the average over the four time points and used it to remove the sloppy global normalization constant: 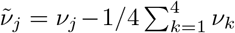. Let *C* be the resulting covariance matrix after this linear transformation, assuming each *ν_j_* has an independent estimated experimental standard deviation of 0.4. The corresponding normalized values of the model are denoted by 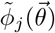, calculated from the log2 ratios of Flag amount over Myc amount at N-1, 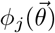 (see Method Section H3 histone exchange model), in the same way. We then used

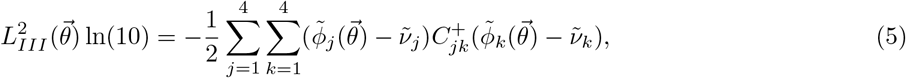

where *C*^+^ is the Moore-Penrose inverse (pseudoinverse) of *C*.

### H3 histone exchange model

To obtain the Flag and Myc amounts in a given model with given parameter values and then determine 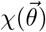 and 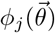, we used the histone pool and nucleosome turnover models in Dion et al., 2007 and assumed that the Myc H3 and Flag H3 amounts in the histone pool are given by

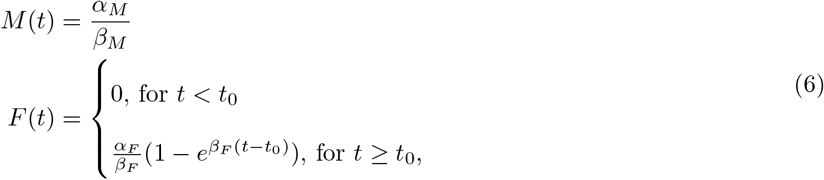

where we used the production rates *α_F_* = 50 / min, *α_M_* = 10 / min, the degradation rates *β_F_* = 0.01 / min, *β_M_* = 0.03 / min and the lag time *t*_0_ = 15 min which were fitted in Dion et al., 2007.

For *t* > *t*_0_, the probability that a newly assembled nucleosome contains a Flag H3 is given by

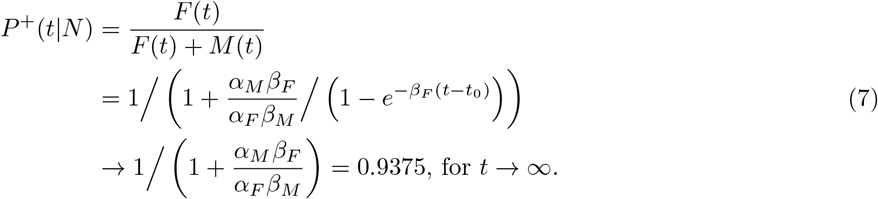

In Dion et al., 2007, the conditional probability that a given nucleosome at site *l* at time *t* contains a Flag H3 then fulfills the ordinary differential equation

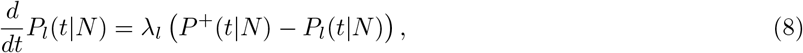

with *λ_l_* being an effective turnover rate.

In our case, the dynamics of the three promoter nucleosomes is coupled, determined by the transition rate matrix 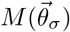 of a given on-off-slide model. If we include different nucleosome types (i.e. Flag and Myc) into the model, the eight promoter configurations are replaced by all 27 possibilities to arrange no, a Flag or a Myc nucleosome at each of the three sites. Each assembly reaction rate in the extended Flag/Myc transition rate matrix 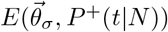 is given by the normal assembly rate, where Flag or Myc nucleosomes are just normal nucleosomes, in 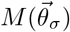 times either *P*^+^(*t*|*N*) or 1 − *P*^+^(*t*|*N*), for a new Flag or Myc nucleosome, respectively. The rates of sliding and disassembly of Flag or Myc nucleosomes are equal to the normal sliding and disassembly rates. The solution 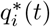 of

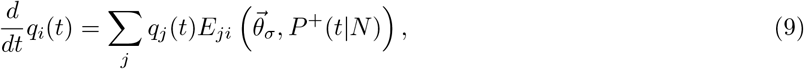

is the probability of extended configuration *i* at time *t*, where *σ* is fixed in the repressed state and *i* and *j* now denote the 27 extended promoter states. The log2 ratios of Flag at N-1 over Flag at N-2 amount, 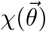, and Flag over Myc amounts at N-1, 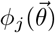, of each model then correspond to log2 ratios of sums of 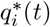 over suitable configurations *i* at the corresponding time points with Flag or Myc nucleosomes at the wanted sites.

### Sensitivity analysis

In order to determine how sloppy the found best parameter values for a given model are, we performed a simple sensitivity analysis, by calculating the log10 likelihood *L_I_* + *L_II_* + *L_III_* along certain directions from the best fit point in logarithmic parameter space. We found that an approximation of the real likelihood function by a second order Taylor expansion at the best fit point worked only in a small area, as expected in a highly non-linear setting, but too small to determine parameter sloppiness properly.

As a compromise between properly scanning the parameter space and computational feasibility, we chose a small number of test directions: each fitted parameter value individually, the eigenvectors of the numerical Hessian of the likelihood function at the best fit, as well as the numerical gradient, which can be non-zero if the best fit point lies on the boundary. We ignored the boundary during the sensitivity analysis, to also take into account sloppiness that “reaches over” the boundary. Along these directions we tested in exponentially increasing steps from the best fit position which positions in parameter space lead to a decrease of the likelihood by ≈ 50%, i.e. a log10 likelihood ratio change of ≈ 0.30, which is of similar order as the log10 likelihood differences within our group of satisfactory models. We then obtained “error bars” for each parameter by taking the largest deviation of the log10 parameter value at the 50% likelihood level from the best value found in all tested directions (Table S2 and Table S3).

### Effective chromatin opening and closing rates

The effective trajectory in time of the regulated process rate from the value of the repressed state to the value of the activated state depends on how fast the cell senses the phosphate starvation and subsequent signal processes. To obtain a reasonable upper bound for the chromatin opening rate, we assumed the regulation happens instantaneously, i.e. the activated rate value of the regulated process applies immediately at the change of the medium for a population in repressed state. Then the promoter configuration distribution decays exponentially towards the activated steady state with a rate well approximated by the negative eigenvalue of the transition rate matrix closest (but not equal) to zero, taking into account the fitted time scale. This “effective chromatin opening rate” is an upper bound of how fast a given model can switch to the activated state. Conversely, we did the same calculations for the “effective chromatin closing rate”, which is an upper bound of how fast a given model can switch to the repressed state.

### Sticky N-3 experiments

Strains “Periodicity 1” and “Periodicity 2” used for restriction enzyme accessibility assays were generated by transformation of linear fragments of plasmids ECS53 and ECS56, respectively, into the wild type strain BY4741 as described in Small et al., 2014. For the Periodicity 1 mutant, the sequence GTTTTCTCATGTAAGCGGACGTCGTC inside the *PHO5* promoter was replaced with GTTTTCTTATGTAAGCTTACGTCGTC. For Periodicity 2, GCGCAAATATGTCAACGTATTTGGAAG was replaced with GCGCAAATATGTCAAAGTATTTGGAAG. Strains were grown in YPDA medium to logarithmic phase for repressive (+Pi) and shifted from logarithmic phase to phosphate-free YNB medium (Formedia) over night for inducing (-Pi) conditions. Nuclei preparation, restriction enzyme digestion, DNA purification, secondary digest, agarose gel electrophoresis, Southern blotting, hybridization and Phosphorimager analysis were as in Musladin et al., 2014. Secondary digest was with HaeIII for both ClaI and HhaI digests probing N-2 or N-3, respectively. The probe for both ClaI and HhaI digests corresponded to the ApaI-BamHI restriction fragment upstream of N-3.

## Acknowledgments

We thank Eliza Small (Northwestern University Feinberg School of Medicine, Chicago, USA) for providing the plasmids ECS53 and ECS56.

Furthermore we thank Oliver J. Rando and his group for providing the fitted nucleosome pool parameters used in Dion et al., 2007 and Amine Nourani and his group for providing the Flag-H3 MNase-ChIP values for the N-1 and N-2 sites of the two replicates in Rufiange et al., 2007.

This work was funded by the German Research Foundation (Deutsche Forschungsgemeinschaft, DFG) via the Collaborative Research Clusters SFB863 (to UG) and SFB1064 (to PK). Michael Wolff is a member of the Graduate School of Quantitative Biosciences Munich (QBM).

## Supplementary Material

**Table S1:**
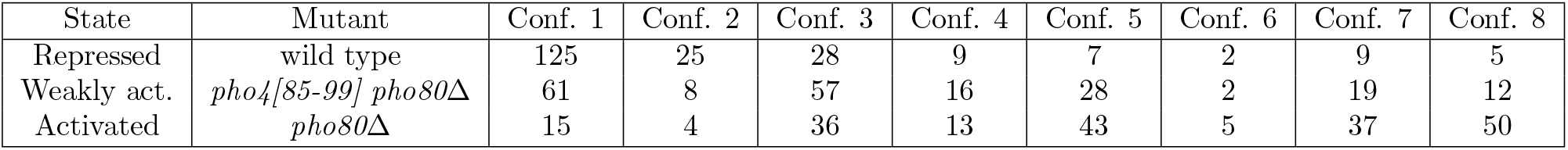
Number of occurrences of each of the eight promoter nucleosome configurations from electron microscopy of single *PHO5* gene molecules in Brown et al., 2013. All cells are grown in high-phosphate conditions which leads to a repressed *PHO5* promoter in wild-type. *pho80* knockout mutants simulate phosphate starvation and exhibit the activated *PHO5* promoter state.

**Table S2:**
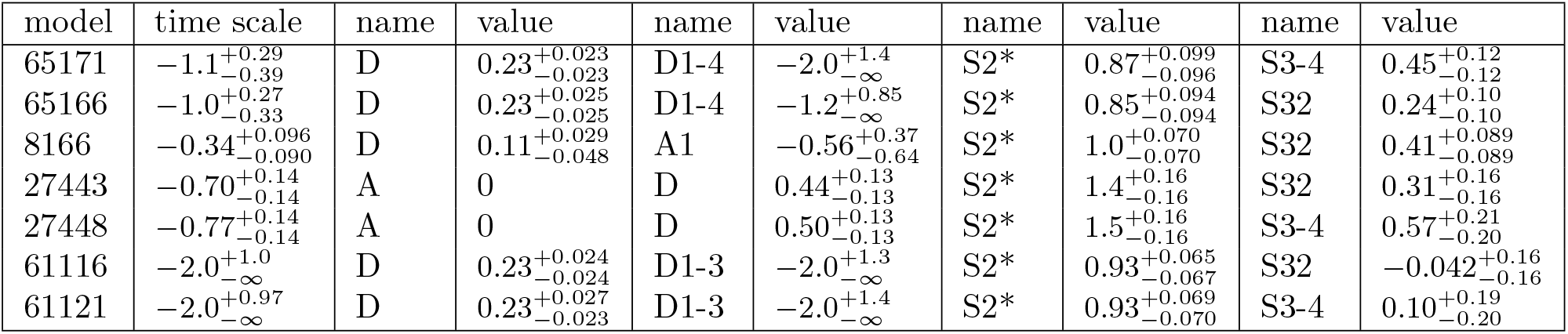
Rate values of constitutive processes (name with relative value on log10 scale) for each of the satisfactory models after the combined fit to all data sets in stage 3 shown in Figure 4. The time scale is given in 1 h^−1^ and also on log10 scale and needs to be added to each individual relative rate value. We also tested the sensitivity of our models with respect to rate changes in certain directions in parameter space (see Methods). ± values for each process correspond to the highest found change from the best process rate value in all tested parameter directions which lead to a ≈ 50% decrease in likelihood and indicate the sloppiness of a rate value (—∞ means the log10 rate value could be made arbitrarily small).

**Table S3:** Rate values of regulated processes (name with value on log10 scale) for each of the satisfactory models after the combined fit to all data sets in stage 3 shown in Figure 4. Same as Table S2, but for regulated instead of constitutive processes.

| model | time scale | name | rep. value | weakly act. value | act. value |
| --- | --- | --- | --- | --- | --- |
| 65171 | $-1.1^{+0.29}_{-0.39}$ | A | $0.80^{+0.051}_{-0.050}$ | $0.41^{+0.045}_{-0.040}$ | 0 |
| 65166 | $-1.0^{+0.27}_{-0.33}$ | A | $0.80^{+0.051}_{-0.049}$ | $0.41^{+0.041}_{-0.040}$ | 0 |
| 8166 | $-0.34^{+0.096}_{-0.090}$ | A | $0.94^{+0.060}_{-0.058}$ | $0.46^{+0.045}_{-0.045}$ | 0 |
| 27443 | $-0.70^{+0.14}_{-0.14}$ | A2 | $1.4^{+0.14}_{-0.14}$ | $0.94^{+0.15}_{-0.15}$ | $0.43^{+0.19}_{-0.19}$ |
| 27448 | $-0.77^{+0.14}_{-0.14}$ | A2 | $1.5^{+0.14}_{-0.14}$ | $1.0^{+0.15}_{-0.15}$ | $0.51^{+0.18}_{-0.18}$ |
| 61116 | $-2.0^{+1.0}_{-\infty}$ | A | $0.79^{+0.048}_{-0.048}$ | $0.41^{+0.041}_{-0.040}$ | 0 |
| 61121 | $-2.0^{+0.97}_{-\infty}$ | A | $0.79^{+0.048}_{-0.048}$ | $0.41^{+0.040}_{-0.040}$ | 0 |

**Figure S1:**
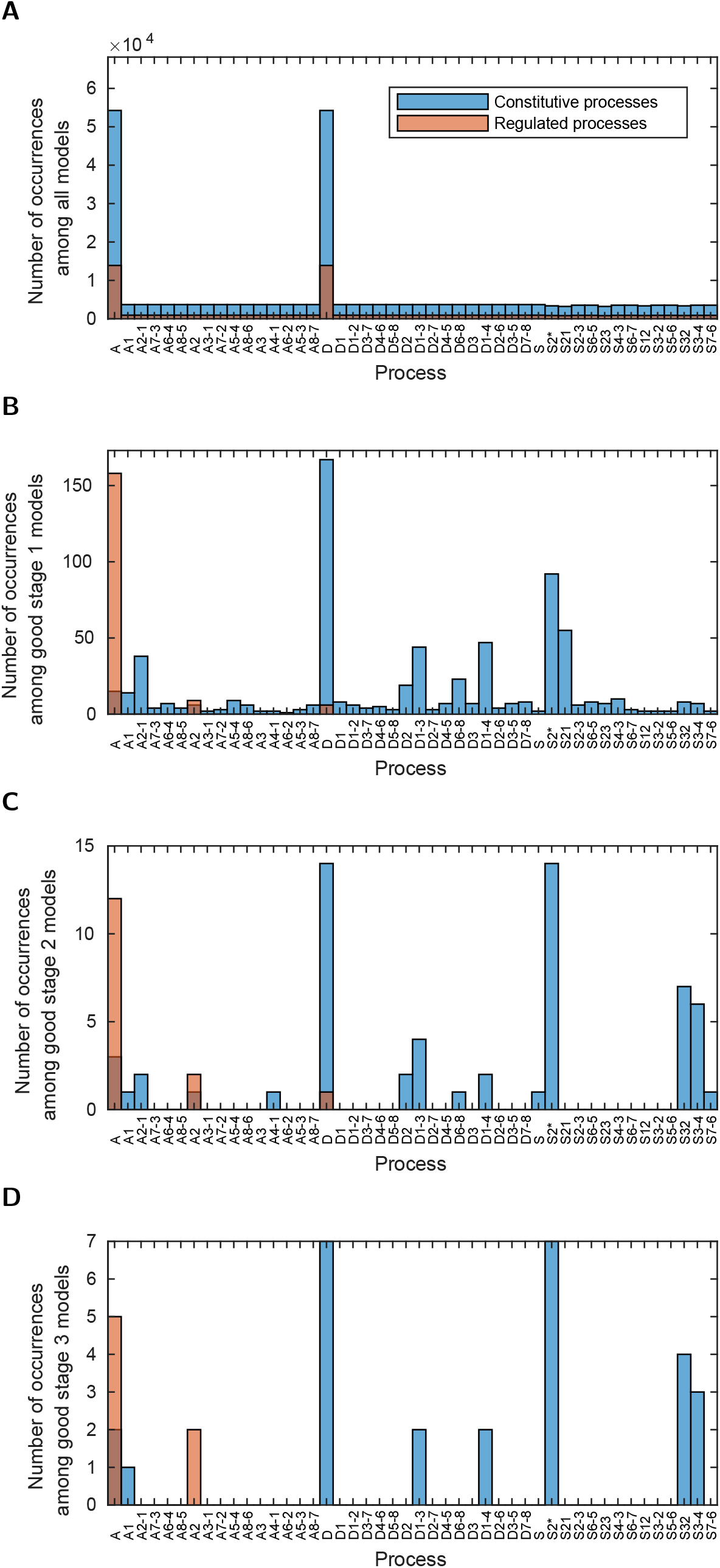
Occurrences of the different processes in the models with satisfactory likelihood at the different stages. (**A**) in all 68 145 analyzed models, (**B**) in all 173 good models of stage 1, (**C**) in all 15 good models of stage 2, (**D**) in all 7 satisfactory models after stage 3. In each plot the y-axis limit is the number of the considered models allowing the comparison of the relative occurrences between the four cases.

**Figure S2:**
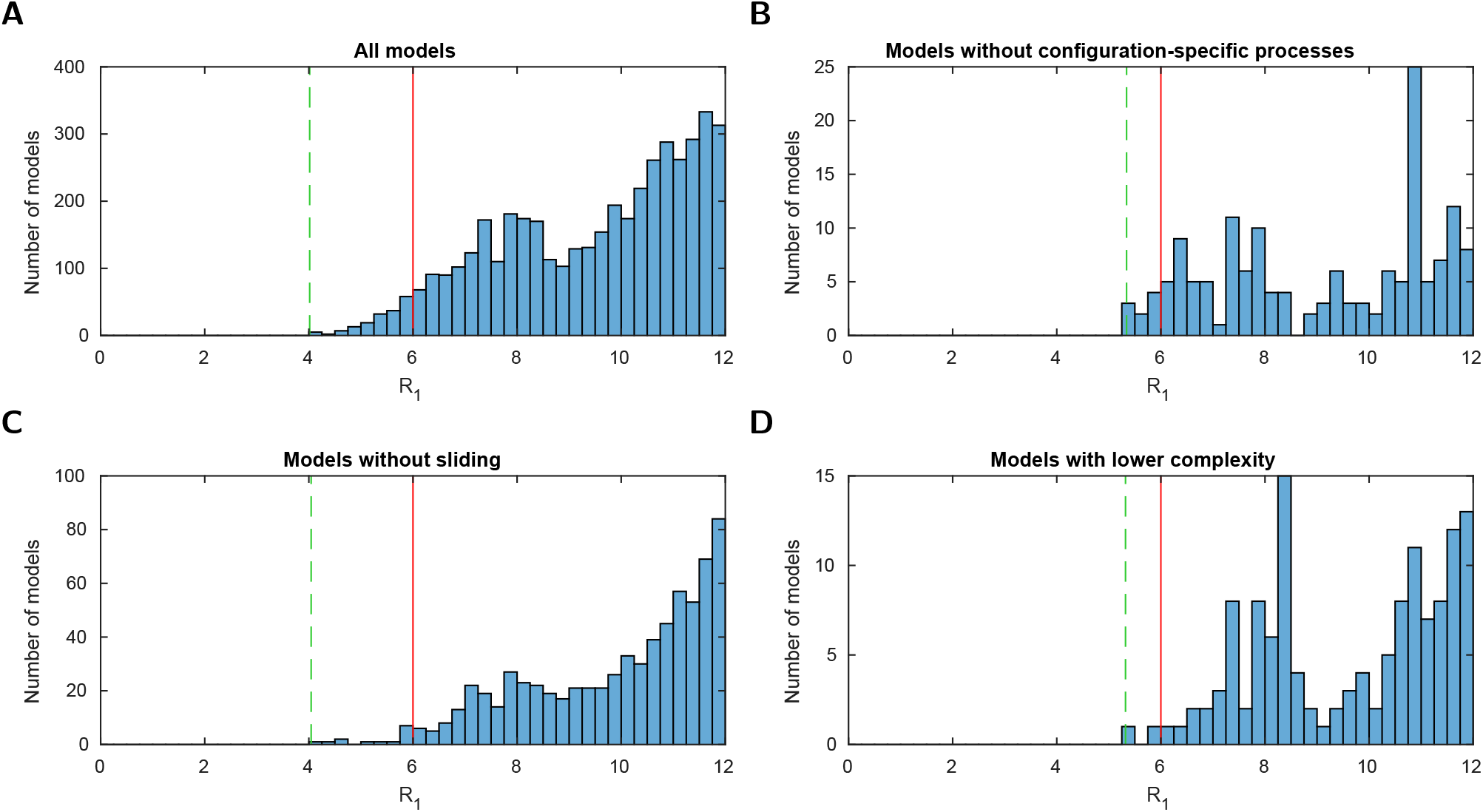
Histograms of the logarithmic likelihood ratio with respect to the perfect fit likelihood in stage 1 (*R*_1_), i.e. using the configurational data of Brown et al., 2013 only. (**A - D**) For all models and three model subsets. 0 on the x-axis corresponds to a perfect fit. Dashed green line: value of the best model (within the subset). Red line: threshold for a satisfactory fit of *R_max_* = 6. Sliding processes are not needed to find agreement with the measured configuration statistics alone. 2 models with lower complexity, i.e. with one fit parameter less, are below the threshold *R_max_*.

**Figure S3:**
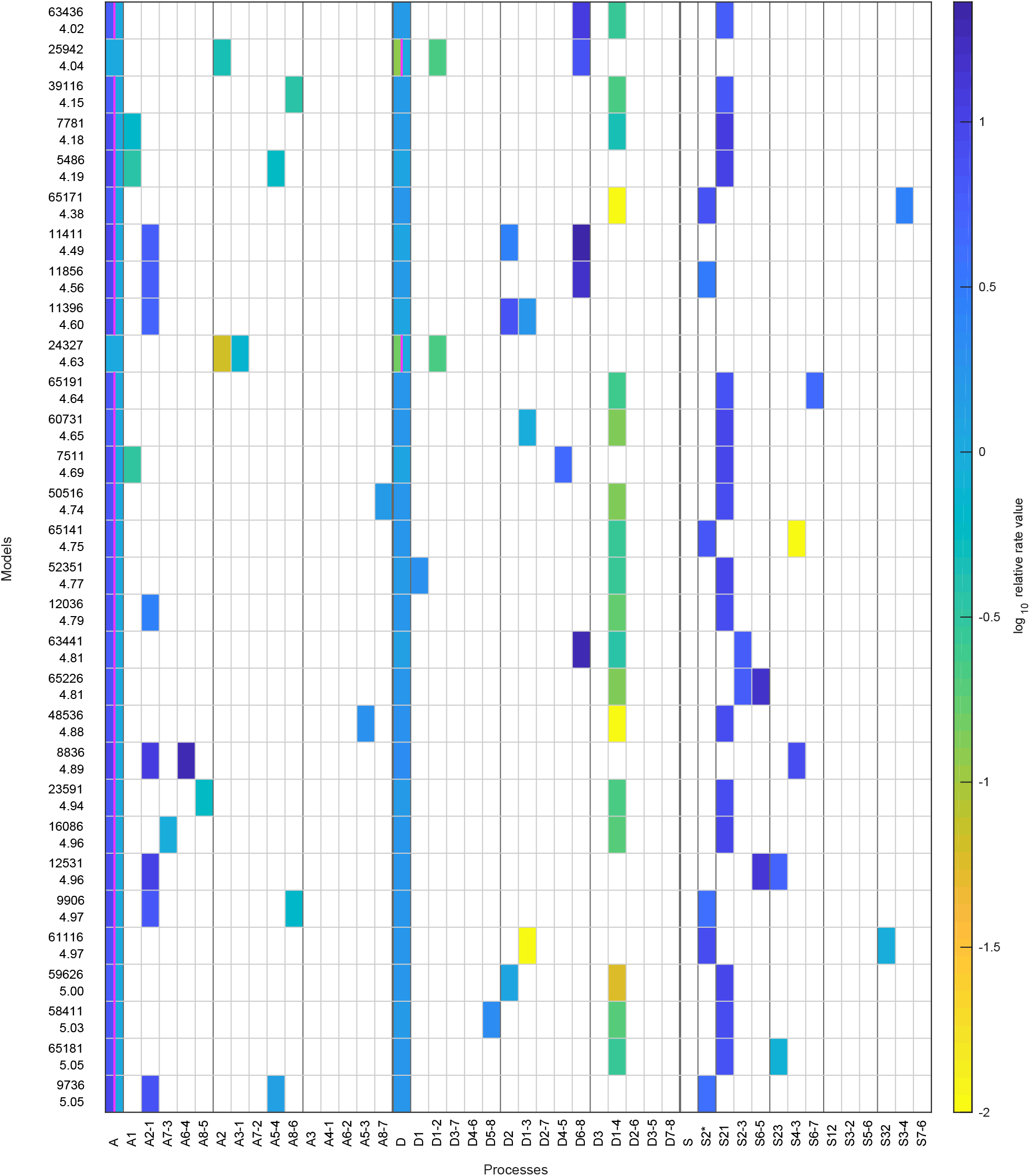
The top 30 models with likelihood above the threshold after the first stage. The colored boxes in each row show the model processes and their rate values. White boxes denote the absence of a process in a model. Regulated processes are separated into two differently colored boxes for repressed (left half) and activated (right half) promoter state. Weakly activated rate values are not shown here. On the left side are the model number and the log10 ratio of the best possible likelihood and the model likelihood, *R*_1_.

**Figure S4:**
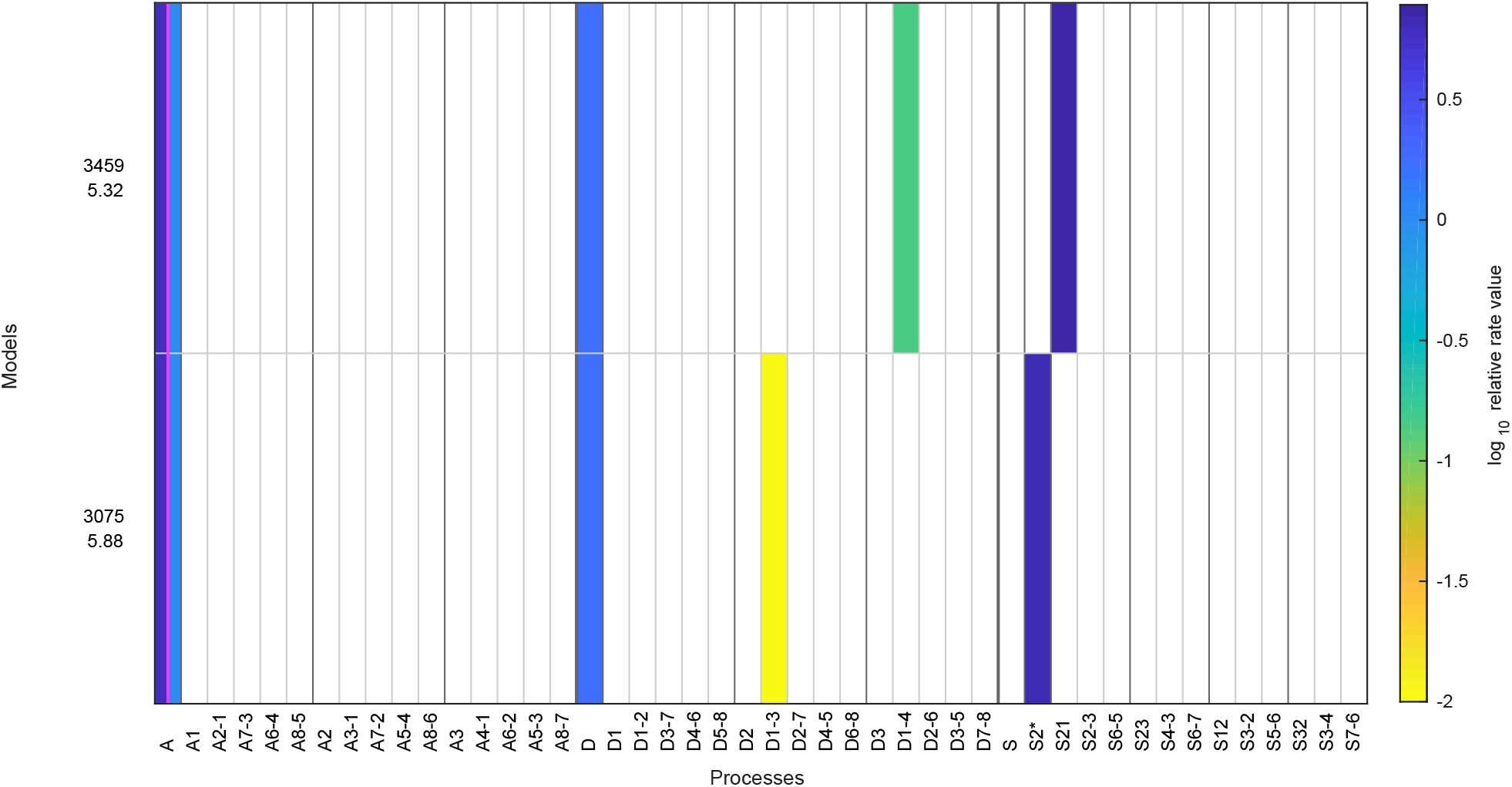
Same as in Figure S3, but showing satisfactory stage 1 regulated on-off-slide models with complexity reduced by one, i.e. models with up to only 6 fit parameters.

**Figure S5:**
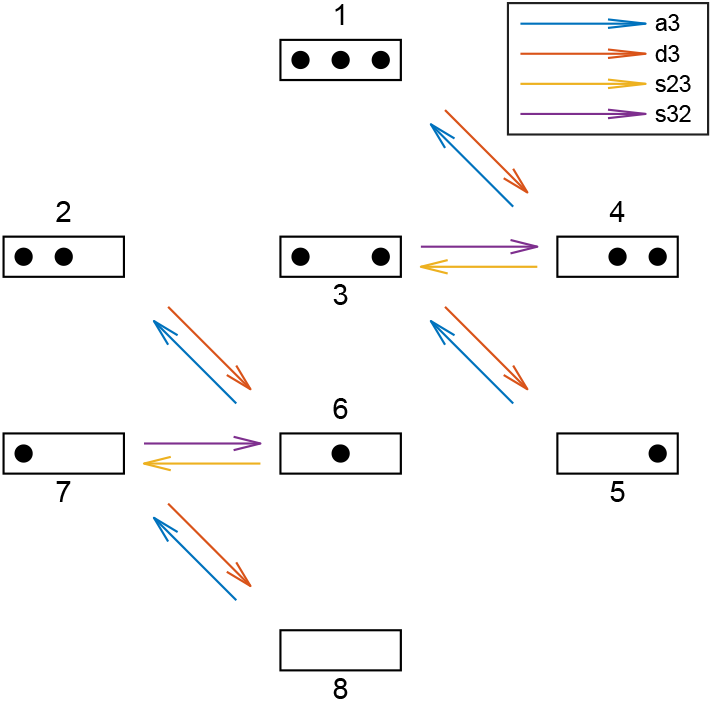
Reactions involving the N-3 position. “a3”, “d3”, “s23” and “s32” denote sets of reactions. For each model the reaction rates are governed by the model’s processes as in stage 1. Simultaneously, to test models for the agreement with the sticky N-3 mutation experiments, the model’s reaction rates of assembly at N-3 (a3), disassembly at N-3 (d3), sliding from N-2 to N-3 (s23) and sliding from N-3 to N-2 (s32), each obtain a prefactor whose values are found by maximizing the combined likelihood of the configurational data and the sticky N-3 mutant accessibility fold-changes (see Methods).

**Figure S6:**
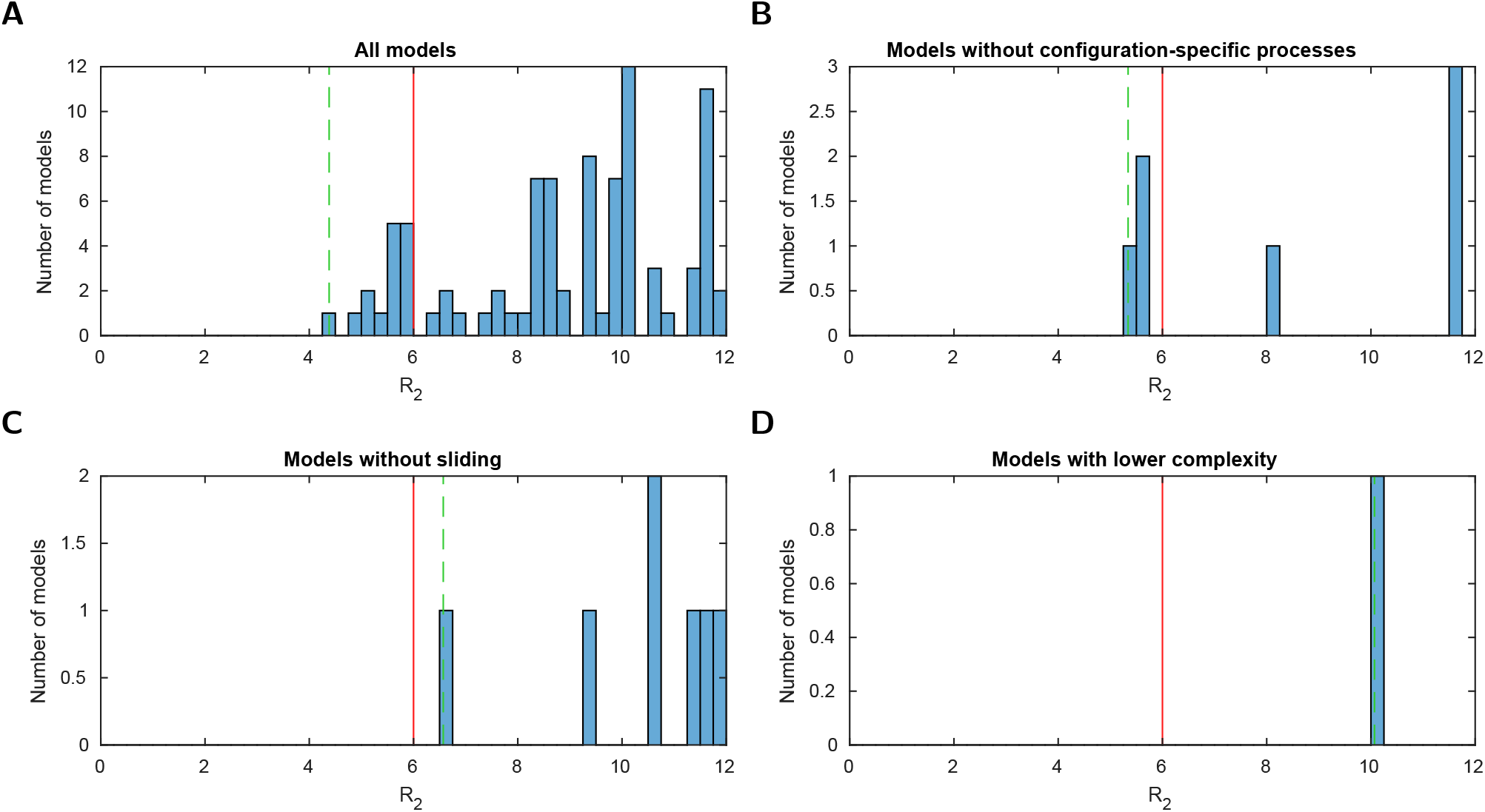
Histograms of the logarithmic likelihood ratio with respect to the perfect fit likelihood in stage 2, *R*_2_, i.e. using the configurational data of Brown et al., 2013 and the sticky N-3 accessibility data (Table 1). (**A - D**) For all models and three model subsets. 0 on the x-axis corresponds to a perfect fit. Dashed green line: value of the best model (within the subset). Red line: threshold for a “good” fit of *R_max_* = 6.

**Figure S7:**
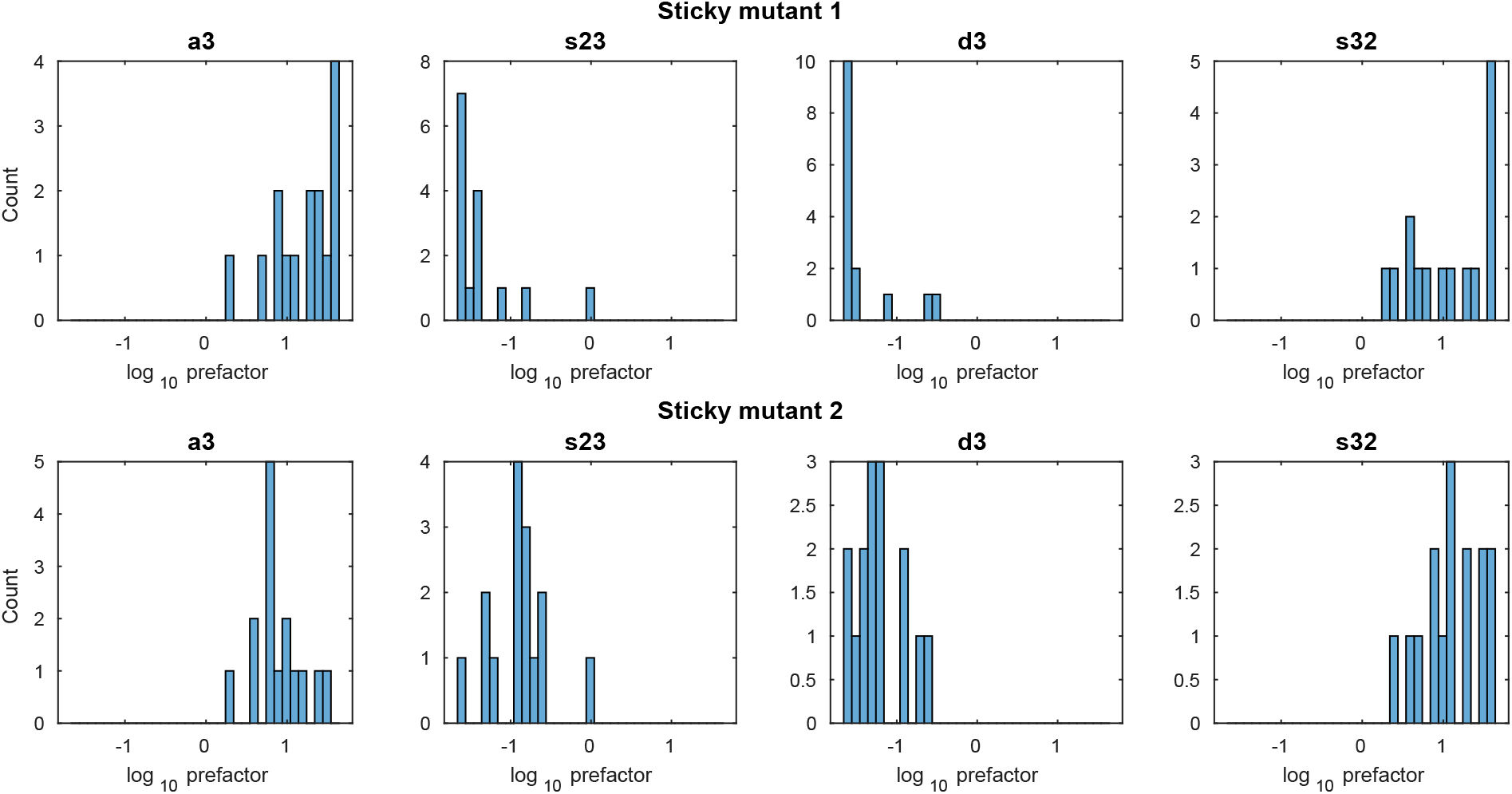
Histograms of the four prefactor values for the reaction sets a3, s23, d3 and s32 for both sticky mutants and all 15 models with maximum likelihood above the threshold in stage 2.

**Figure S8:**
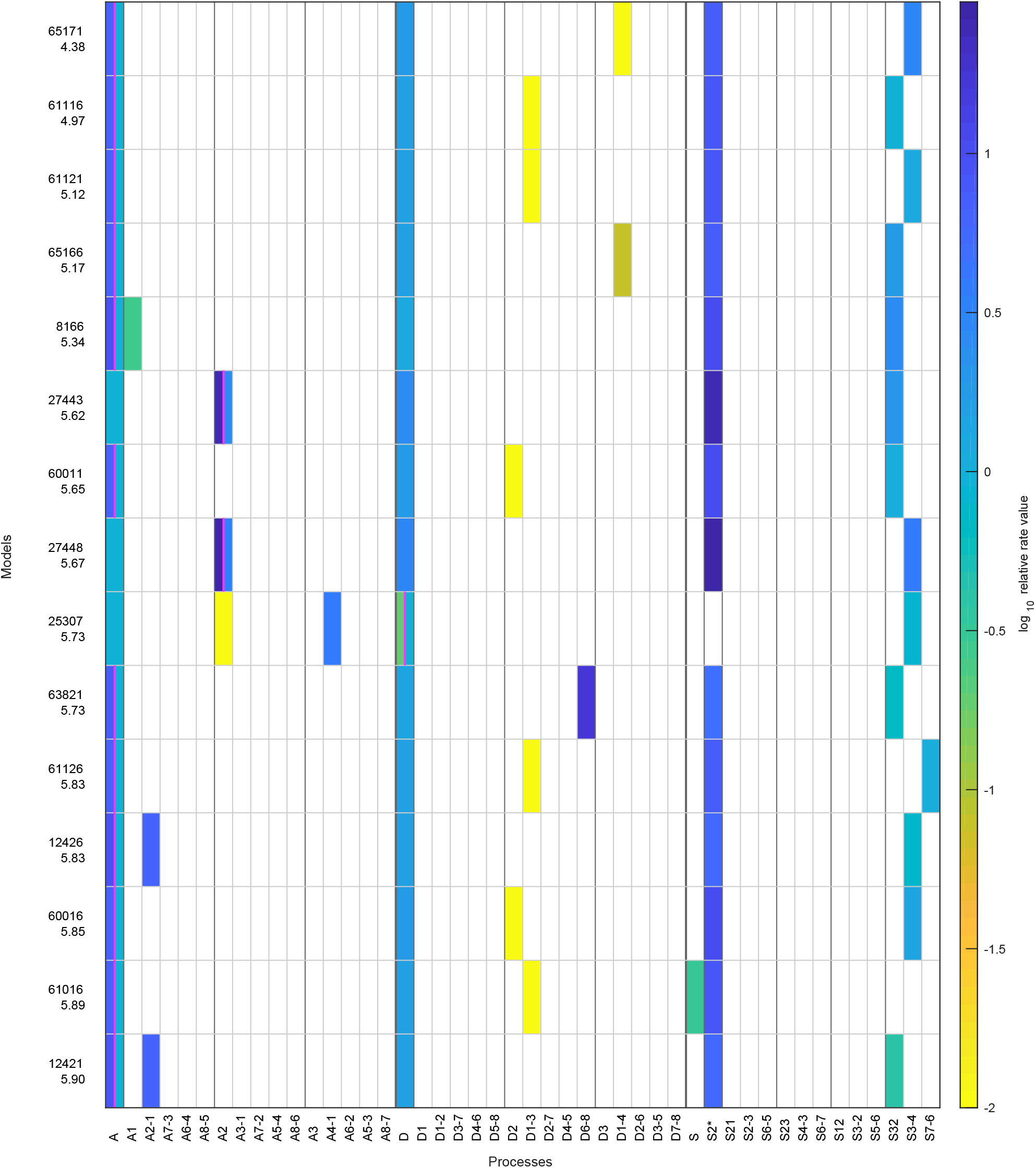
Same as in Figure S4, but showing the regulated on-off-slide models with likelihood above the threshold with up to 7 parameters after stage 2, with the stage 2 log10 likelihood ratios, *R*_2_, written below the model numbers.

**Figure S9:**
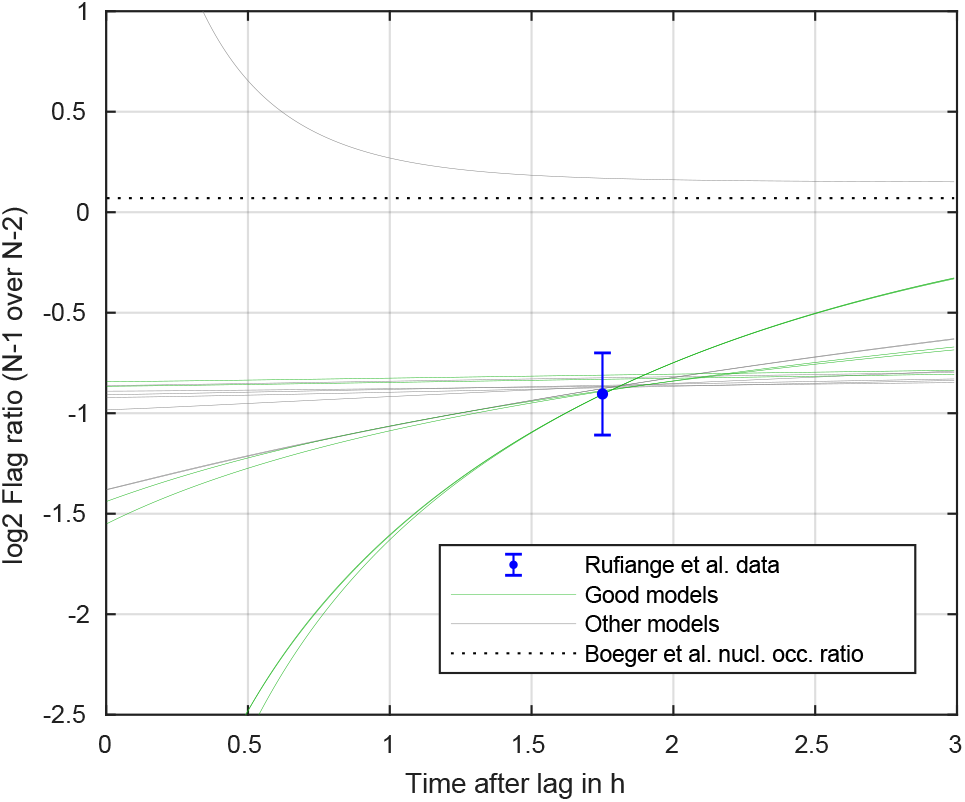
Flag amount ratio of N-1 over N-2, calculated for all models with satisfactory likelihood after stage 2. Green line shows the satisfactory models of stage 3. The dashed line shows the steady state ratio that eventually all models should reach closely. The three groups of green lines match the three model groups of the main text and Figure 4: almost constant lines (group 3), slightly rising lines (group 1) and quickly rising lines (group 2).

**Figure S10:**
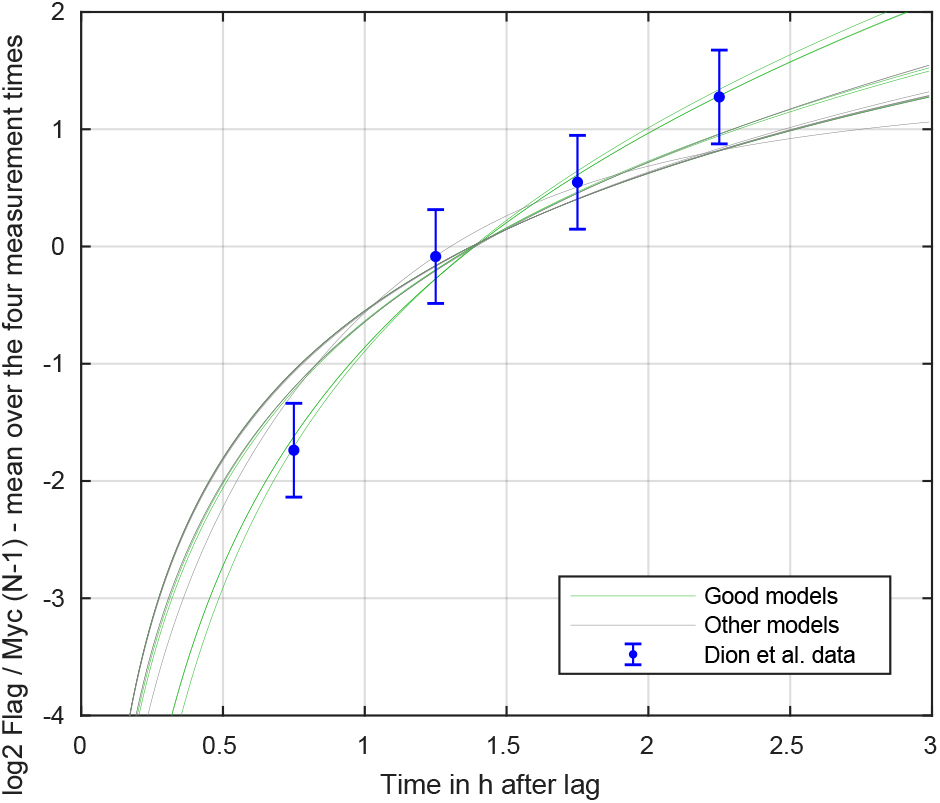
Shifted Flag over Myc amount ratio at N-1 position, calculated for all models with satisfactory likelihood after stage 2. Green line shows the satisfactory models of stage 3.

**Figure S11:**
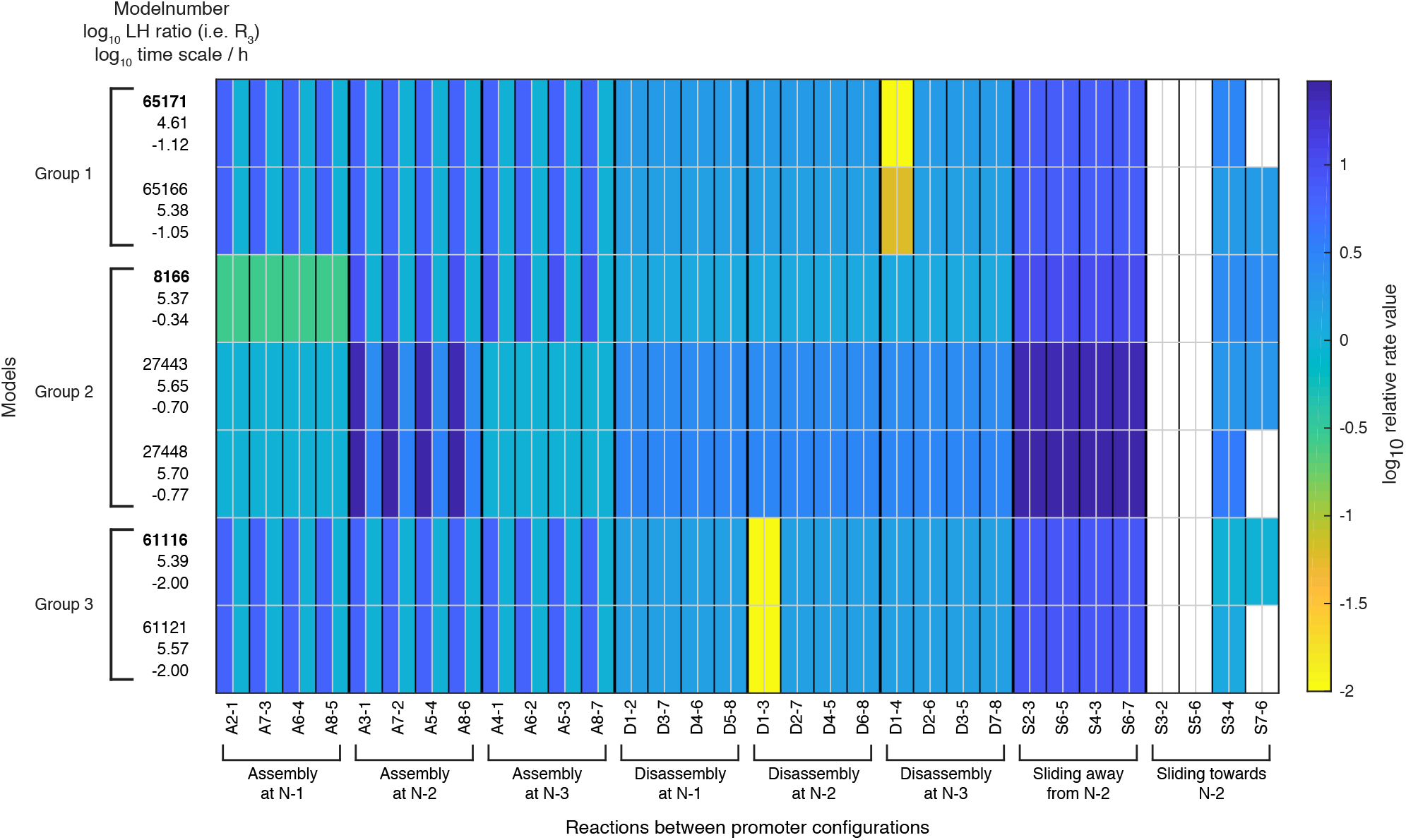
Rate values for each of the 32 reactions when using the processes and their rate values of each model as shown in Figure 4. The colored boxes show the relative rate values with respect to the global assembly process rate in activated state. White fields correspond to a reaction rate of zero. For a given reaction, the left half shows the repressed value, the right half the activated value. Weakly activated rate values are not shown. The left column shows for each model: the model number, the log10 likelihood ratio *R*_3_ and the log10 time scale parameter value.

**Figure S12:**
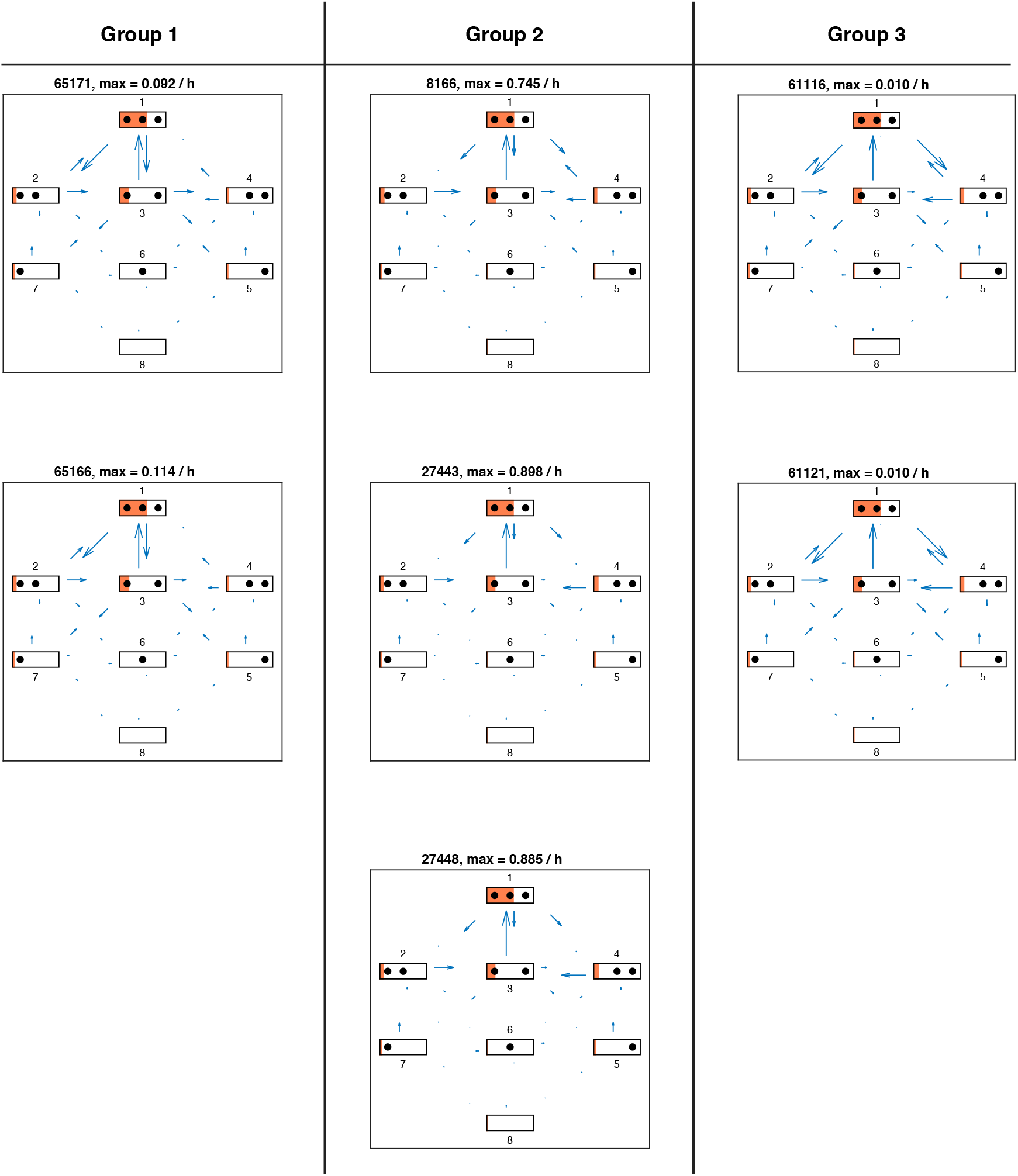
Directional fluxes in repressed promoter state for each satisfactory model in stage 3. The length of the flux arrows indicates the amount of net flux with respect to the maximum value for each model stated above. The orange filling of each state symbol shows the steady state probabilities. The models are grouped with respect to similarities in the site-centric net fluxes (Figure S16).

**Figure S13:**
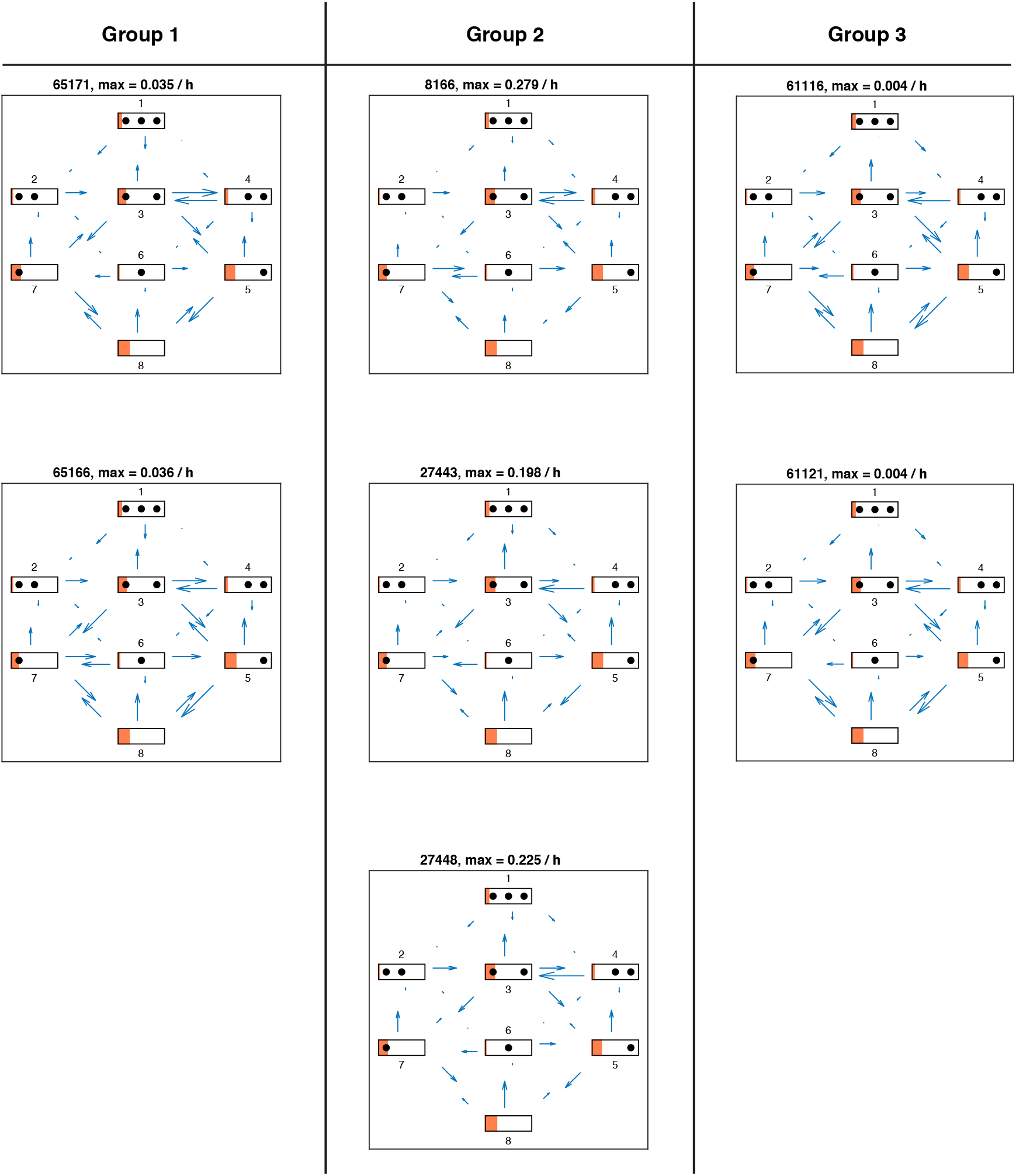
Directional fluxes in activated promoter state for each satisfactory model in stage 3.

**Figure S14:**
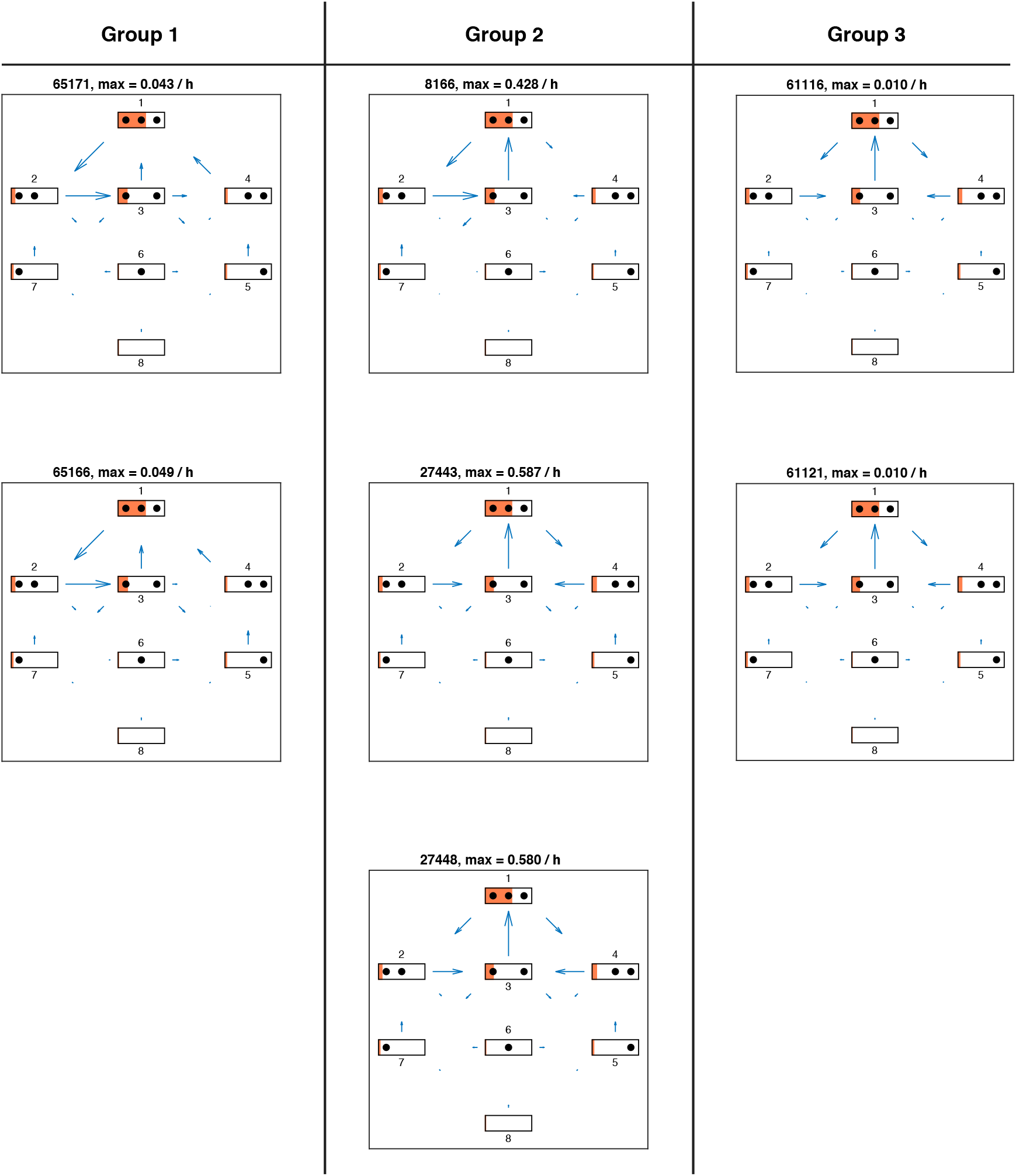
Net fluxes in repressed promoter state for each satisfactory model in stage 3. The length of the flux arrows indicates the amount of net flux with respect to the maximum value for each model stated above. The orange filling of each state symbol shows the steady state probabilities. The models are grouped with respect to similarities in the site-centric net fluxes (Figure S16).

**Figure S15:**
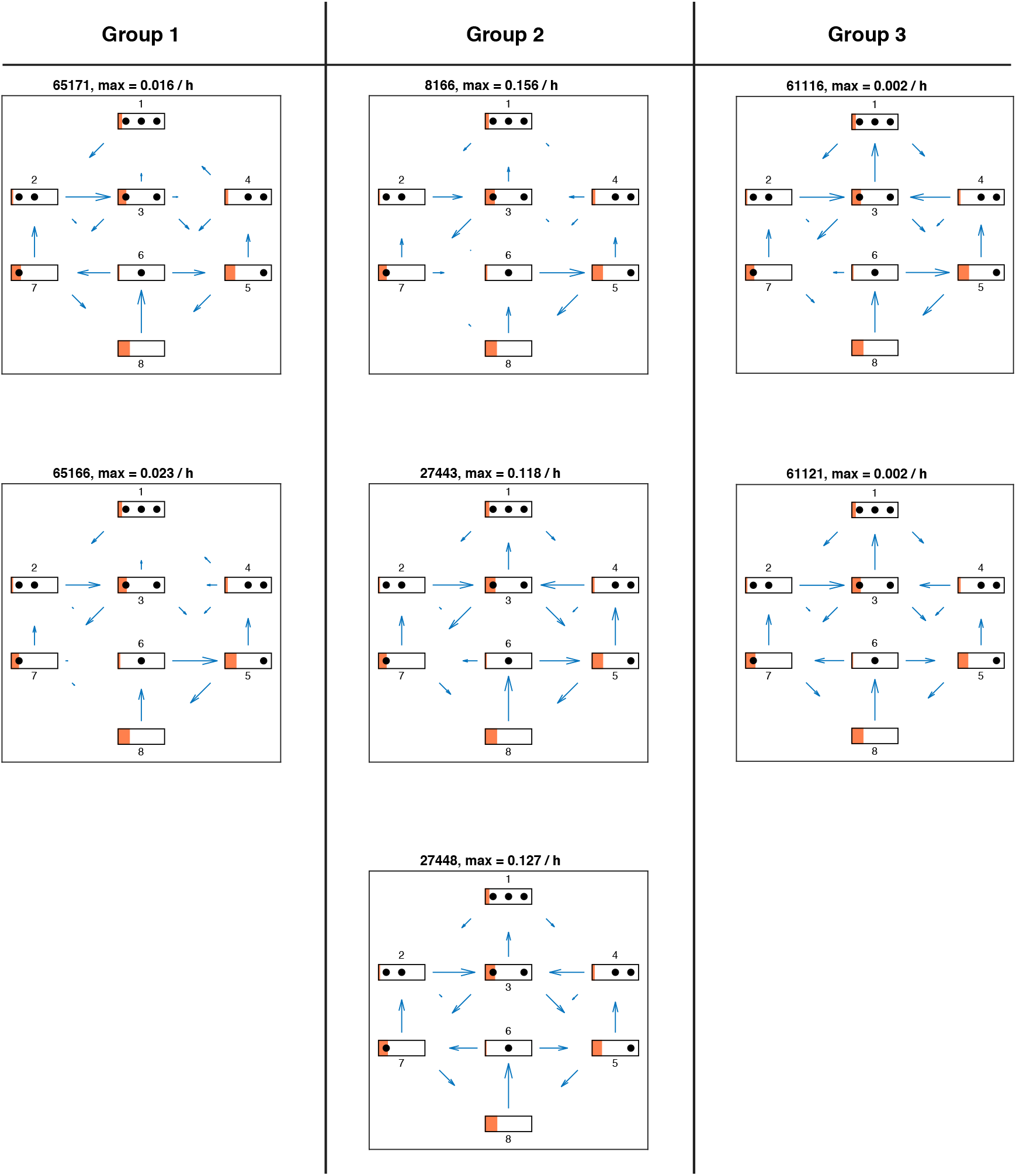
Net fluxes in activated promoter state for each satisfactory model in stage 3.

**Figure S16:**
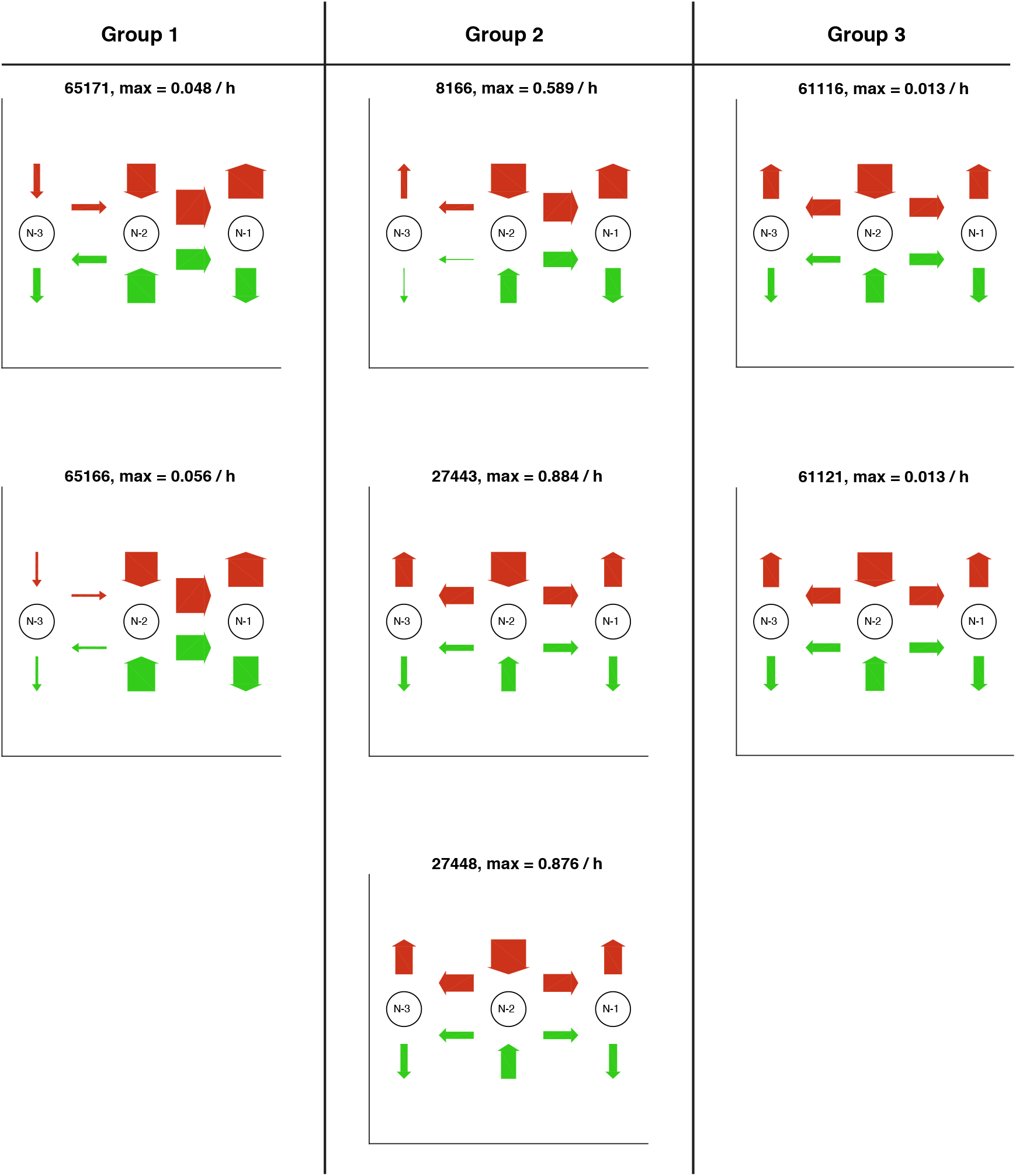
Site-centric net fluxes in active (green) and repressed (red) promoter state for each satisfactory model in stage 3. Obtained by summing all assembly/disassembly net fluxes at each site and sliding net fluxes between N-1 and N-2 as well as N-2 and N-3. The arrow thickness indicates the amount of flux with the maximum value stated above. The models are grouped with respect to similarities in the site-centric net fluxes (also see main text).

## Notes

### Competing Interest Statement

The authors have declared no competing interest.

